# Fatty Acid Metabolism and The Oxidative Stress Response Support Bacterial Predation

**DOI:** 10.1101/2023.12.11.571100

**Authors:** Rikesh Jain, Nguyen-Hung Le, Lionel Bertaux, Jean Baudry, Jérôme Bibette, Yann Denis, Bianca H. Habermann, Tâm Mignot

**Affiliations:** Aix-Marseille Université - CNRS UMR 7283, Institut de Microbiologie de la Méditerranée and Turing Center for Living Systems, Marseille, France; Aix-Marseille Université - CNRS UMR 7283, Institut de Microbiologie de la Méditerranée, Marseille, France; Laboratoire Colloïdes et Matériaux Divisés, Institut Chimie, Biologie, Innovation, UMR 8231, ESPCI Paris, CNRS, Université Paris Sciences et Lettres, 75005 Paris, France; Aix-Marseille Université – CNRS FR3479, Institut de Microbiologie de la Méditerranée, Marseille, France; Aix Marseille University, CNRS, IBDM UMR 7288, Turing Center for Living Systems, Marseille, France

## Abstract

Despite growing awareness of their importance in soil ecology, the genetic and physiological traits of bacterial predators are still relatively poorly understood. In the course of a *Myxococcus xanthus* predator evolution experiment, we discovered a class of genotypes leading to enhanced predation against diverse species. RNA-seq analysis demonstrated that this phenotype is linked to the constitutive activation of a predation-specific program. Functional analysis of the mutations accumulated across the evolutionary time in a two-component system and Acyl-CoA-manipulating enzymes revealed the critical roles of fatty acid metabolism and antioxidant gene induction. The former likely adapts the predator to metabolites derived from the prey while the latter protects predatory cells from reactive oxygen species (ROS) generated by prey cells under stress and released upon lysis during predation. These findings reveal interesting parallels between bacterial predator-prey dynamics and pathogen-host cell interactions.

**Significance Statement:** This study illuminates the largely unexplored genetic and metabolic strategies used by bacterial predators in soil ecosystems. Through experimental evolution in *Myxococcus xanthus*, we discovered that more efficient predators accumulate mutations that activate a genetic program for predation. This program simultaneously triggers a metabolic shift favoring fatty acid degradation for energy production and upregulates antioxidant gene expression, enhancing protection against reactive oxygen species generated during prey cell lysis. This adaptive mechanism proves advantageous across a wide range of prey species, suggesting that metabolic adaptation plays a crucial role in the evolutionary trajectory of bacterial predators within their natural ecological niche.

## Introduction

Intensive agriculture and the systematic use of chemicals over decades of practice have deprived soil of microbial biodiversity, greatly endangering food chain productivity in the relatively short term(1, 2). Restoring these ecosystems could pave the way for more sustainable practices, which is especially urgent given that this problem is aggravated by global warming(3). Bacterial predators have recently come to the forefront for their role in shaping microbial soil ecosystems(4, 5). Among them, the *Myxobacteria* act potentially as key taxa within the microbial food web(4, 6, 7). Characterized as generalist predators, *Myxobacteria* feed on a vast array of microorganisms, including diverse bacterial and fungal species(8). Their predatory nature offers promising avenues in combating plant infections. Notably, certain species of *Myxobacteria* have been shown to efficiently counteract infections from various pests, spanning from fungi to pathogenic bacteria(9, 10). Therefore, understanding how these predators interact with their prey and digest them could help determine and leverage their function in soil ecology.

Predation in *Myxobacteria* is a complex interplay of various factors, from motility and secretion of secondary metabolites to specialized mechanisms of contact-dependent killing. We and others recently discovered that when *Myxococcus xanthus* cells establish contact with the prey cells, they temporarily halt motility and activate two unique cellular machineries – a suspected Tad pilus-like system (Kil) and an unconventional Type-3 secretion system (T3SS)(11, 12). These systems act sequentially to trigger prey cell plasmolysis (Kil), and subsequently lyse the dead prey cells (T3SS) to obtain the nutrients that fuel *Myxococcus* growth(12). On the chromosome, the *kil* and *t3ss* systems are encoded by three prominent genetic clusters, collectively comprising over 60 predicted genes(11–13). This large genetic repertoire is likely associated with the high versatility of the Kil/T3SS system, which can detect and kill a diverse array of monoderm and diderm bacterial species of distinct evolutionary origins and ecologies (action against fungi has not been tested(9, 14)).

Contact-dependent killing is only the first step of the process. Downstream, several metabolic adaptations also contribute in the predation process. *Myxococcus* must also metabolize prey- derived nutrients and overcome potential prey defenses. This was suggested by transcriptomic studies encompassing diverse prey species, which consistently unveiled substantial shifts in the genetic expression profiles(15–17). In addition to the *kil* genes, lytic enzymes and genes encoding secondary metabolites, metabolic genes encoding iron siderophore pathways and fatty acid degradation are systematically induced(11, 15, 16).

To explore the physiological changes that occur during predation and what genetic pathways are required during such an atypical growth phase, we conducted experimental evolution selecting for enhanced predation by the *Myxococcus* DZ2 laboratory strain. The rationale was that by identifying the traits that evolve to improve predation, we might gain insight into previously unknown mechanisms of predation and predator-prey interactions. We discovered that evolved predators turn on a predation-specific metabolic program constitutively. Analyzing this genetic switch, we find that enhanced predation requires two distinct adaptations, a metabolic shift towards fatty acid consumption and the resistance to reactive oxygen species (ROS) liberated from prey lysis.

## Results

### Conserved *Myxococcus* transcriptional response across different prey

To determine if there is a core genetic response to predation, we sequenced the *Myxococcus* transcriptome during interactions with three different prey species—*Escherichia coli*, *Bacillus subtilis*, and *Caulobacter crescentus*—under uniform experimental conditions (see Methods).

Echoing previous findings(15, 16), *Myxococcus* demonstrated significant changes in gene expression (Wald test, *p.adj* < 0.05) in response to each prey: 2165 genes with *E. coli*, 2379 genes with *B. subtilis*, and 3575 genes with *C. crescentus* (Figure 1A-1C, Table S1). While overall gene expression profiles showed parallels between interactions with *E. coli* and *B. subtilis*, they differed markedly when *Myxococcus* encountered *C. crescentus* (Figure S1). The underlying mechanisms for this disparity warrant further investigation.

**FIGURE 1:**
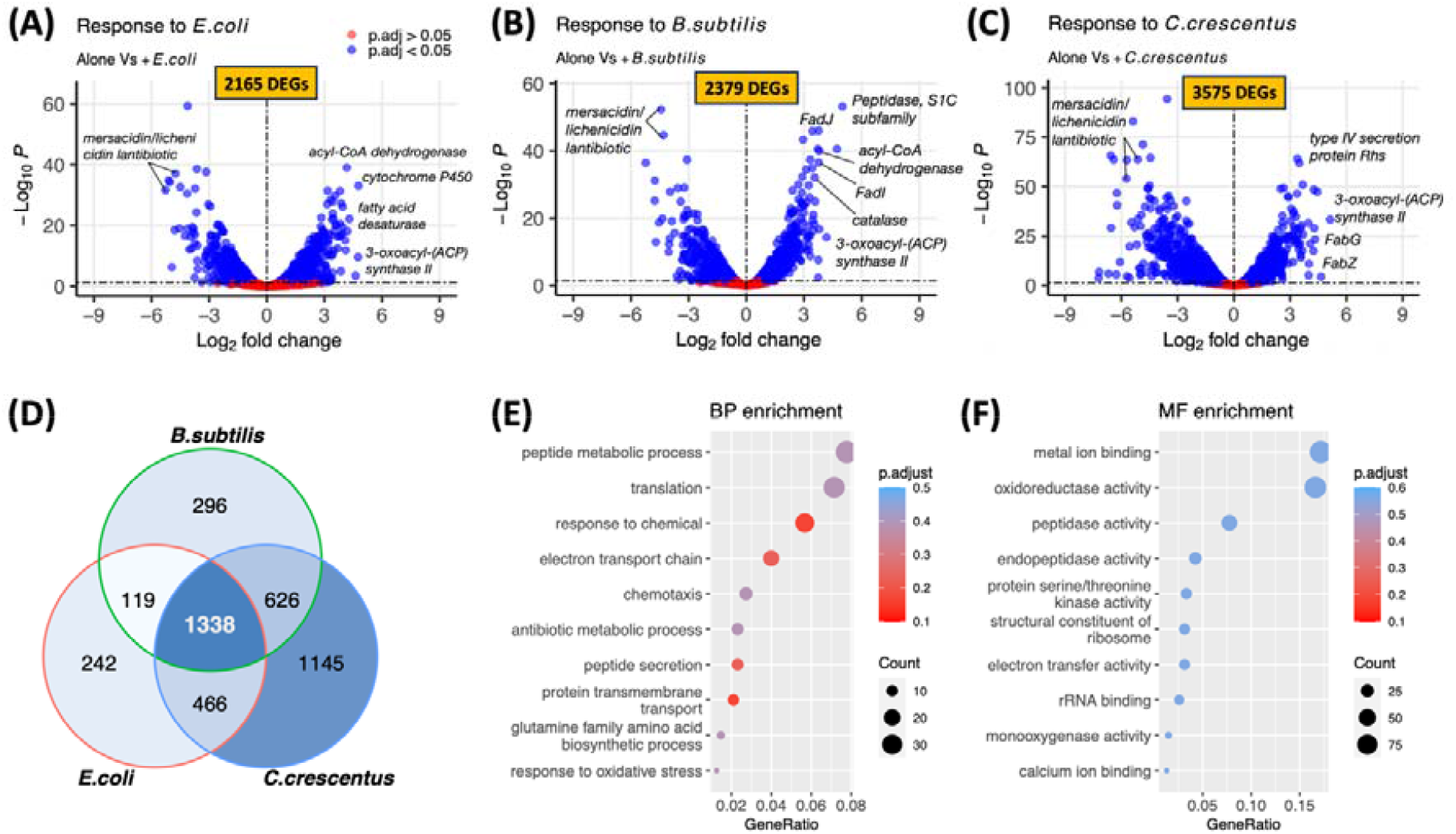
Transcriptomic shifts in wild-type *M. xanthus* in response to varied prey. (**A)** Contrasting gene expression patterns of *M. xanthus* between solitary state and *E. coli* predation (RNA-seq, n=3 biological replicates). Genes with a positive log2 fold change value are upregulated in the *M. xanthus* + prey condition, while genes with a negative log2 fold change value are up-regulated in the solitary condition. DEGs stands for Differentially Expressed Genes (*p.adj* < 0.05, Wald test). **(B)** Gene expression variations in *M. xanthus* during solitary state versus *B. subtilis* predation (n=3). **(C)** Alterations in gene expression of *M. xanthus* during solitary state versus *C. crescentus* predation (n=3). **(D)** Common and distinct DEGs in *M. xanthus* predation across three distinct prey types. **(E)** Gene ontology term enrichment for biological processes (BP) in common genes linked to predation on all three prey. **(F)** Gene ontology term enrichment for molecular functions (MF) in common genes involved in predation on all three prey.

We subsequently focused on gene responses that were consistent across all three prey species (Figure 1D). 1338 *M. xanthus* genes (Table S2) exhibited differential expression upon *M. xanthus* encounters with any prey. Remarkably, out of these 1338 genes, a staggering 1329 (>99%) showed a consistent pattern, either upregulation or downregulation, across all three comparisons. To gain further insights, we conducted Gene Ontology (GO) analysis based on the current *Myxococcus* genome annotation, with a focus on enrichment in two domains: Biological Processes (BP) and Molecular Function (MF). GO analysis revealed enrichment of the functions related to translation, respiration, metabolism, and amino acid biosynthesis (particularly Glutamine), suggesting that *Myxococcus* undergoes active growth when exposed to these prey species (See Figures 1E and 1F). In line with previous studies, we also observed an enrichment of genes associated with secondary metabolite production and degradative enzymes, such as proteases and peptidases (Figures 1E and 1F). Intriguingly, both regulatory and chemotaxis systems, especially the Ser/Thr kinases, exhibited changes (Figures 1E and 1F). While the biological functionality of the majority of *M. xanthus* Ser/Thr kinase systems (encompassing over 100 modules) remain unclear(18), it is possible that many are predominantly operational during the predation phase. Lastly, we noted that genes linked to the oxidative stress response were ubiquitously enriched across all three interactions.

### Enhanced predatory capabilities of *Myxococcus* variant on *E. coli* prey

RNA-seq provided insights into physiological patterns related to predation. To identify which of these adaptations are crucial, we next aimed to identify gain-of-function variants that exhibit improved predatory efficiency.

To this end, we employed *E. coli* as prey. Plating a *Myxococcus* colony adjacent to an *E. coli* colony on a CF medium (detailed in Methods) results in total predation of the prey within 72 hours (Figure 2A). However, when the medium is supplemented with glucose, a sugar that *E. coli* can consume but not *Myxococcus*(19), prey growth is observed and predation is not as efficient (Figure 2A). In fact, predation is completely inhibited at glucose concentrations exceeding 0.1%. This property allowed for an evolution experiment to select superior predatory phenotypes by increasing the concentration of glucose in the media, thereby enhancing prey resistance through evolutionary cycles (Figure 2B). We thus inoculated *Myxococcus* at the center of an *E. coli* colony and allowed a full predatory cycle before transferring to a fresh *E. coli* plate for a new cycle, gradually increasing glucose concentration every 5 cycles (Figure 2B). After 20 cycles and approximately 600 generations of *M. xanthus* (approx. 5 months), we isolated one clone (M1) capable of killing *E. coli* at 0.1% glucose concentration (Figure S2). Glucose uptake in *E. coli* results in acetate release and consequently, medium acidification. Considering that *Myxococcus* can adapt to acid stress via the secretion of diffusible molecules(20), we ascertained that the M1 phenotype was not a consequence of such adaptation, confirming the superior performance of the M1 over the WT in a buffered medium (Figure S3).

**FIGURE 2:**
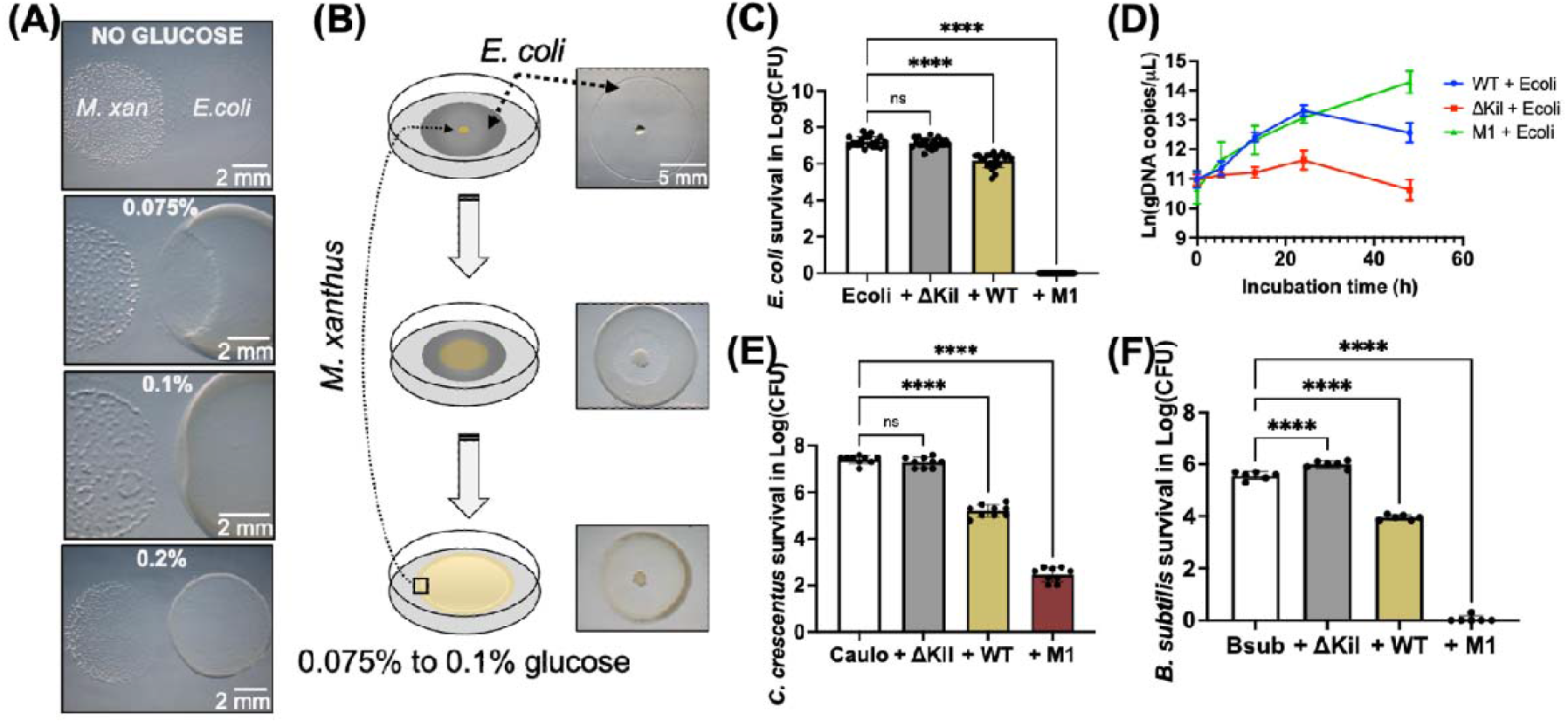
Evolution of a super predator from WT *M. xanthus*. **(A)** In a co-incubation experiment lasting 72 hours, *M. xanthus* and *E. coli* colonies were spotted near to each other on CF media with varying glucose concentrations (ranging from 0% to 0.2%). The efficiency of *M. xanthus* predation decreases with increasing glucose concentrations. **(B)** During the evolution experiment, *M. xanthus* cells were placed in the center of an *E. coli* colony. As the *M. xanthus* cells consumed a significant portion of the *E. coli* population and reached the prey colony edge, they were transferred to the center of a new *E. coli* colony. After every 5 cycles, the glucose concentration in the media was gradually raised by 10%. **(C)** Killing assay of *E. coli* MG1655 prey using *M. xanthus* DZ2 WT, predation-deficient ΔKil, and evolved mutant M1 as predators. The bar graph shows the mean prey survival in log10(CFU) after a 24-hour coincubation +/- standard deviation (SD) derived from a minimum of 10 biological replicates. Statistical significance was determined using one-way ANOVA, with post-hoc multiple comparisons performed via the Dunnett test; ns: not significant, **** *p*< 0.0001. **(D)** Growth curves of *M. xanthus* DZ2 WT, predation-deficient ΔKil, and evolved mutant M1 during predation on *E. coli* prey. The scatter plot depicts Ln(gDNA copies/μL) across various predation time points. gDNA concentrations were directly quantified using digital PCR (dPCR). Error bars indicate the SD derived from 3 biological replicates. Comparison of WT and M1 predation efficiency on *B. subtilis* **(E)** and *C. crescentus* **(F)**, assessed by counting prey CFU after predation. The bar graph shows the mean prey survival in log10(CFU) after a 24-hour coincubation +/- SD derived from 3 biological replicates. Statistical significance was determined using one-way ANOVA, with post-hoc multiple comparisons performed via the Dunnett test; ns: not significant, **** *p*< 0.0001.

To rule out any effect that would be linked to glucose adaptation rather than changes in predatory traits, we compared the killing capacity of the M1 and WT strains at a glucose concentration that is permissive for WT predation (0.075%). Counting *E. coli* colony-forming units (CFUs) after 24 hours of co-incubation with *Myxococcus* (see Methods), revealed that while the WT only killed ∼90% of the prey colony, the M1 strain eradicated it entirely (Figure 2C). Investigating whether the elevated predatory efficiency of the M1 correlates with growth and fitness enhancement, we directly quantified *Myxococcus* growth on *E. coli* prey using quantitative digital PCR (see Methods). For the first 24 hours, the WT and M1 variants displayed equivalent growth, but M1 peaked at a higher level by 48 hours, implying a physiological fitness advantage (Figure 2D). As expected, a Δ*kil* mutant did not grow in these conditions, confirming that this assay scores prey-dependent *Myxococcus* growth. We conclude that the M1 variant has evolved enhanced predatory capacities over its WT ancestor, which we further investigated.

We subsequently assessed whether the enhanced predatory capacity against *E. coli* was a specific adaptation or a more general evolved trait effective against other bacterial preys, *B. subtilis* and *C. crescentus*. The predation efficiency was enhanced against both prey species, although to a lesser extent against *C. crescentus* (Figures 2E and 2F). Thus, the M1 strain contains a selected trait(s) that boosts *Myxococcus* ability to target various prey species.

### Genetic basis of superior predatory phenotypes

We next sought to identify the genetic mutations underlying the M1 phenotype. As previously noted, the M1 strain was derived after 20 serial passages (∼600 generations) in our evolution experiment. In fact, the M1 phenotype emerged from a progenitor clone (C10E9) displaying an intermediate enhanced predatory phenotype (Figure 3A and 3B). This suggested that the C10E9 clone represents an evolutionary intermediate to the emergence of the M1 strain.

**FIGURE 3:**
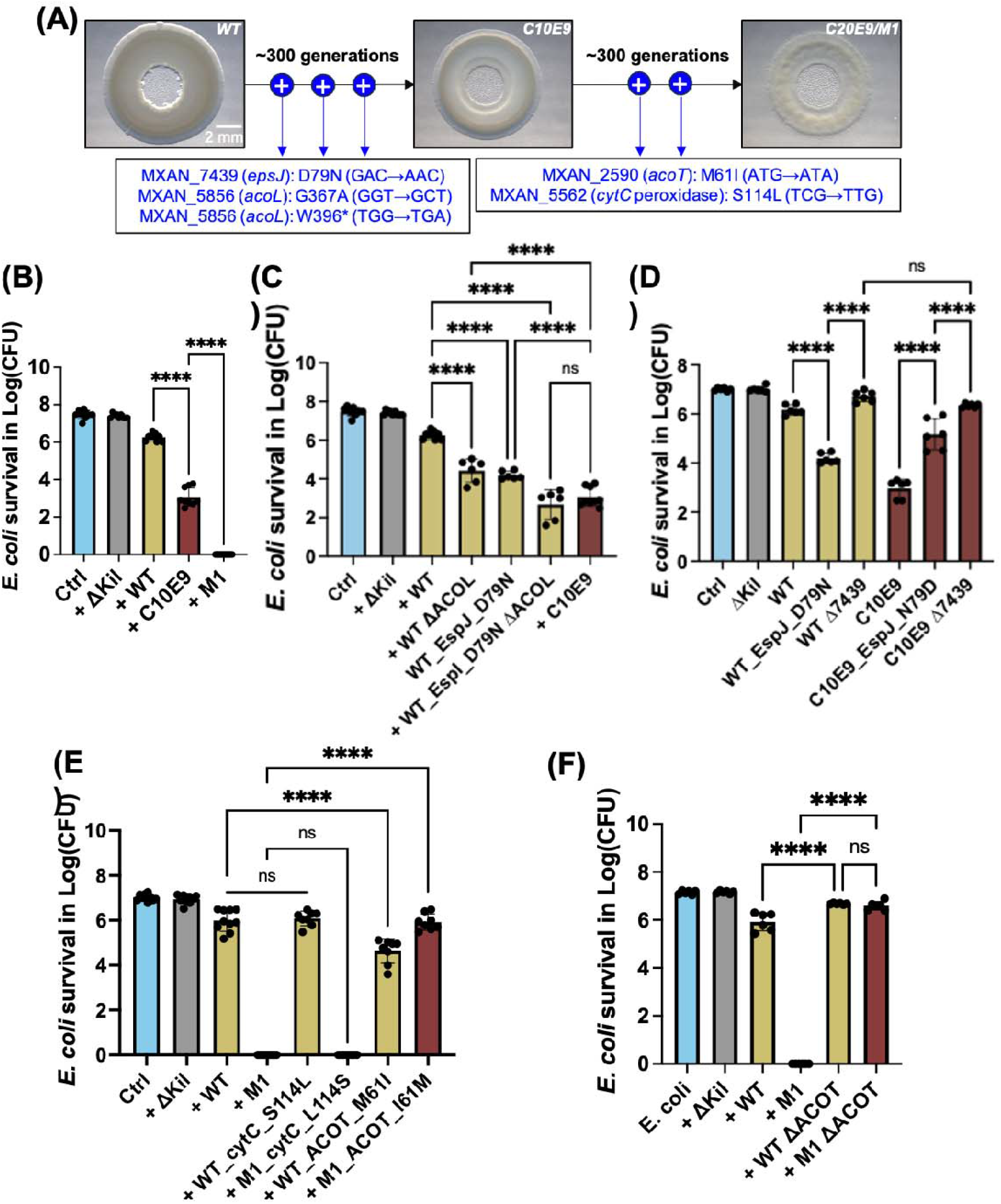
SNPs associated with efficient predation. **(A)** Genome sequencing revealed that the M1(C20E9) strain emerged after selection of an intermediate C10E9 strain. The WT ancestor differs from the C10E9 by 3 SNPs. Two additional SNPs separate the C10E9 transition from the M1. **(B)** Predation efficiency of the evolutionary intermediate C10E9 on *E. coli*. C10E9 exhibits an intermediate predation efficiency, situating between the killing capacities of WT and M1 strains when targeting *E. coli* prey. **(C)** Introduction of D79N mutation in the *epsJ* gene and deletion of MXAN_5856 (*acoL*) separately increases the WT strain predation efficiency. In combination, both mutations increase WT predation to C10E9 level. **(D)** Impact of *epsJ* gene on predation efficiency of WT and C10E9 strains. The reversed mutation N79D of *epsJ* gene in C10E9 strain and *epsJ* gene deletion from both WT and C10E9 strains resulted in diminished predation efficiency, suggesting that the D79N mutation within *epsJ* gene may be conferring a gain-of-function effect. **(E)** *E. coli* prey was subjected to predation by *M. xanthus* WT strains with introduced point mutations: S114L in CytC or M61I in ACOT, as well as M1 strains where these individual point mutations were reverted. **(F)** Impact of ACOT gene deletion on predation efficiency of WT and M1 strains. The deletion of the ACOT gene from both strains resulted in diminished predation efficiency, suggesting that the M61I mutation within the ACOT gene may be conferring a gain-of- function effect. All the bar graphs show the mean *E. coli* survival in log10(CFU) after a 24- hour coincubation with different predator strains +/- SD derived from a minimum of 3 biological replicates. Statistical significance was determined using one-way ANOVA, with post-hoc multiple comparisons performed via the Dunnett test; *ns*: not significant, *** *p*< 0.001, **** *p*< 0.0001.

We sequenced the genomes of both the C10E9 and M1 strains. The C10E9 clone carried three single nucleotide polymorphisms (SNPs): one leading to a D79N alteration in a histidine kinase protein (*epsJ*, *MXAN_7439*) located upstream from the *epsI* response regulator implicated in Exopolysaccharide (EPS) synthesis(21), and two mutations (G367A and W396*) in a gene hypothesized to encode an Acetate-CoA ligase (*acoL, MXAN_5856*). M1 genome sequencing revealed two additional SNPs: one in a gene putatively encoding a cytochrome c peroxidase (*cytC*, *MXAN_5562*) and another in a gene for a putative Acyl-CoA Thioesterase (*acoT*, MXAN_2590, Figure 3A). These mutations led to amino acid changes: a S114L substitution in the cytochrome c peroxidase protein and an M61I substitution in the ACOT protein. This analysis suggests a two-step evolution of the M1 phenotype: initial mutations gave rise to the C10E9 phenotype, which was subsequently enhanced by up to two additional mutations.

To understand the contribution of each mutation in the evolution of the M1 strain, we first explored the mutations that led to the C10E9 phenotype. The D79N mutation in *epsJ* maps to a predicted periplasmic sensor domain of the kinase and may lead to a loss or gain of function (Figure S4A). Conversely, the premature stop codon mutation (W396*) in *MXAN_5856* results in the truncation of its C-terminal domain, encompassing residues 531-609 (Figure S4C). This domain is analogous to the C-terminal region in *Salmonella enterica* Acetate-CoA ligase *acsA* (Figure S4B)(22). In the *Salmonella* enzyme, this domain is essential for Acetyl-CoA synthesis, thus, if *MXAN_5856* indeed encodes an Acetate-CoA Ligase, the W396* mutation likely also leads to its enzymatic inactivation(22).

To identify the function of the mutations, we first introduced the *epsJ^D79N^* mutation and a deletion of *MXAN_5856* (*acoL*) separately and in combination. Each mutation separately increased predation efficiency to an intermediate level between WT and C10E9 (Figure 3C). Notably, the combination of these mutations resulted in the same enhanced predation as the C10E9 strain, demonstrating their combined role in the C10E9 evolution (Figure 3C). Since the deletion of *acoL* captures the effect of the W396* substitution, this mutation leads to a loss of function (and also likely the G367A mutation). To further assess the D79N mutation, we deleted the entire *epsJ* gene in both WT and C10E9 strains, which reduced predation efficiency in both cases (Figure 3D). Reverting the mutation to N79D in the C10E9 strain also reduced its efficiency to a level similar to the single *acoL* mutant (Figure 3D). Thus, the D79N mutation confers a gain of function to the *epsJ* gene. (see further below).

We next examined the importance of the additional mutations in the M1 phenotype. As mentioned above, the additional mutations map in two potential *cytC* and *acoT* genes. Introducing these changes into the WT strain and restoring the WT sequences in the M1 strain showed that the CytC mutation (S114L) did not affect the predatory capabilities of either strain, indicating it is not critical for the M1 phenotype (Figure 3E). In contrast, reintroducing the WT *acoT*^I61M^ allele into the M1 strain almost nullified its predatory advantage (Figure 3F). Thus, the M61I mutation in ACOT is largely responsible for the M1 phenotype. However, reverting the AcoT sequence to WT form did not revert the M1 predatory capability to the C10E9 level, indicating a minor contribution of the CytC mutation to the M1 phenotype (see discussion). Finally, an in-frame deletion of the *acoT* gene in both WT and M1 strains drastically impaired predation (Figure 3D), highlighting the essential role of AcoT in *Myxococcus* predation and demonstrating that the M61I mutation is a gain-of-function mutation.

Altogether the results show that the M1 predatory phenotype required three important mutations: two gain-of-function and one loss of function, demonstrating that the M1 genotype indeed originates from selection for a superior predatory performance. We explore the physiological consequences of these three mutations below.

### Evolved predator strains express a predatory genetic program constitutively

The EpsJ kinase has been shown to phosphorylate the EpsI DNA-binding response regulator *in vitro*, strongly suggesting that its activity activates or modulates transcription of the EpsI/Nla24 regulon, which remains yet poorly defined but is important for EPS synthesis and its association with cyclic-di-GMP(23). Thus, a gain-of-function mutation in EpsJ could be associated with changes in gene regulation in the evolved strains. To test this, we first conducted whole genome transcription analysis of the M1 strain, which revealed that the regulation of up to 2901 genes (Wald test, *p.adj* < 0.05) was altered compared to the WT ancestor (Table S3). Focusing on the 50 most highly up- or down-regulated genes, we observed that many gene expression changes in M1 in the absence of prey, mirrored those seen when WT cells were mixed with any of the three prey species (Figure 4). Specifically, this fraction of the genetic repertoire represented 33% (441 genes) of the gene repertoire normally induced during predation (Table S4). Remarkably, genes involved in oxidative stress response and fatty acids metabolism were constitutively expressed in the M1 strain.

**FIGURE 4:**
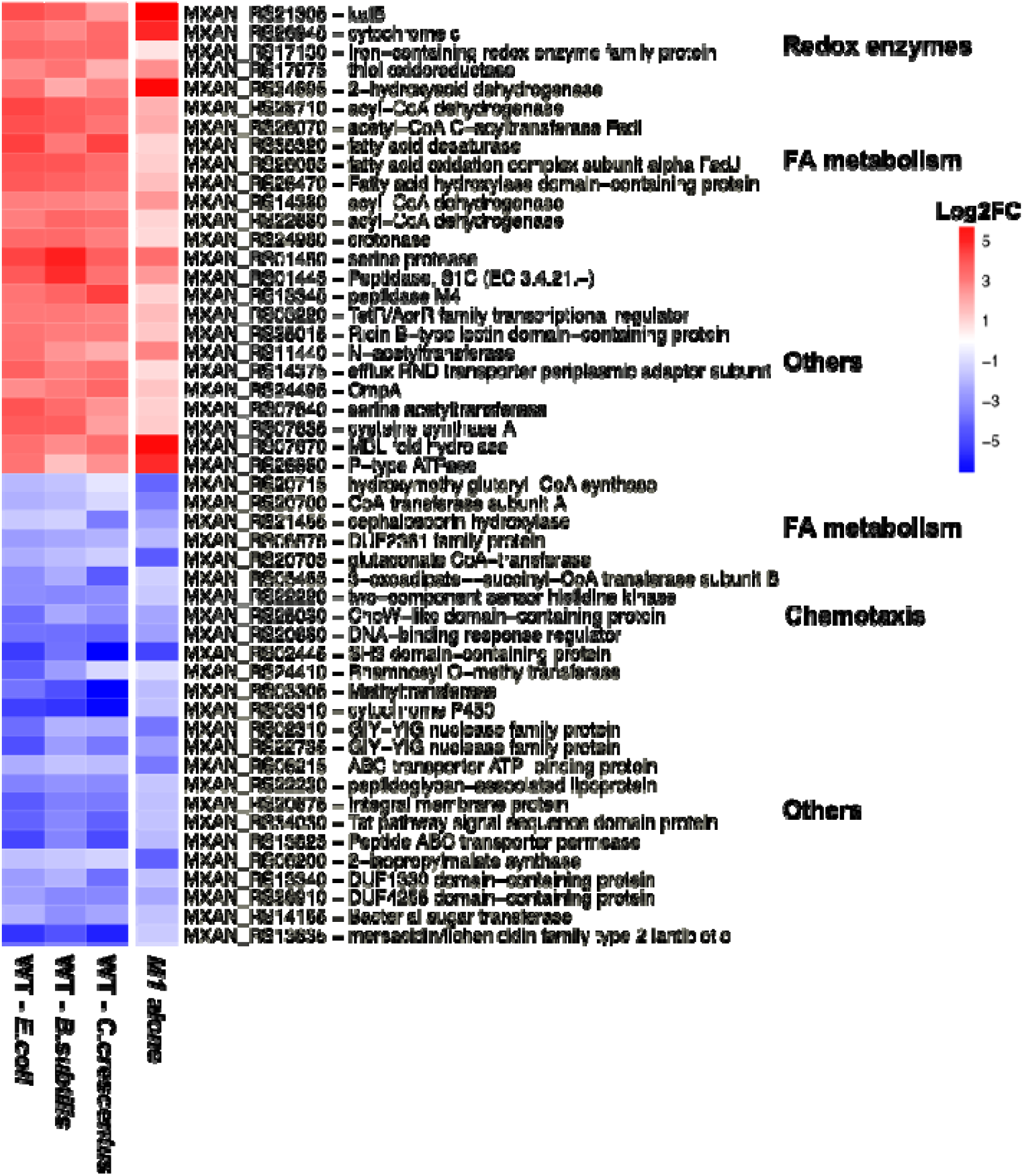
Similar gene expression patterns during WT predation and M1 alone, highlighting predatory readiness in M1. Top 50 genes from the list of 441 genes (Table S4) that show a consistent expression pattern in WT with prey (*E. coli*, *B. subtilis* and *C. crescentus*) as compared to WT alone, and M1-alone as compared to WT alone (RNA-seq, n=3 biological replicates per condition). Notable predation-specific genes expressed in the M1 alone include FA metabolism and redox enzymes like *katB*. The values shown in the heatmaps represent DESeq2normalized counts that have been scaled across conditions for each gene. Each column within the heatmaps represents the mean of 3 independent biological replicates.

### Metabolic shifts observed in the evolved predator strains

Previous reports have shown that WT cells induce the expression of fatty acids β-oxidation pathway genes in the presence of *E. coli* (15, 16). We confirmed this pattern and demonstrated that it also occurs with *B. subtilis* and *C. crescentus* (Figure S5A). To test whether this metabolic program is constitutively induced in the C10E9 and M1 strains, we examined the expression of fatty acid β-oxidation genes. Indeed, we found that both variants constitutively induce these genes (Figures 5A and 5B). To further assess how global predator physiology was impacted along the evolutionary trajectory to the M1 phenotype, we analyzed the expression patterns of genes involved in the TCA cycle and the electron transport chain (ETC) (Figures 5C and 5D). TCA cycle-related genes were upregulated in the C10E9 strain but not in the M1 strain (Figure 5C). Remarkably, ETC genes were downregulated in both evolved strains compared to WT (Figure 5D).

**FIGURE 5:**
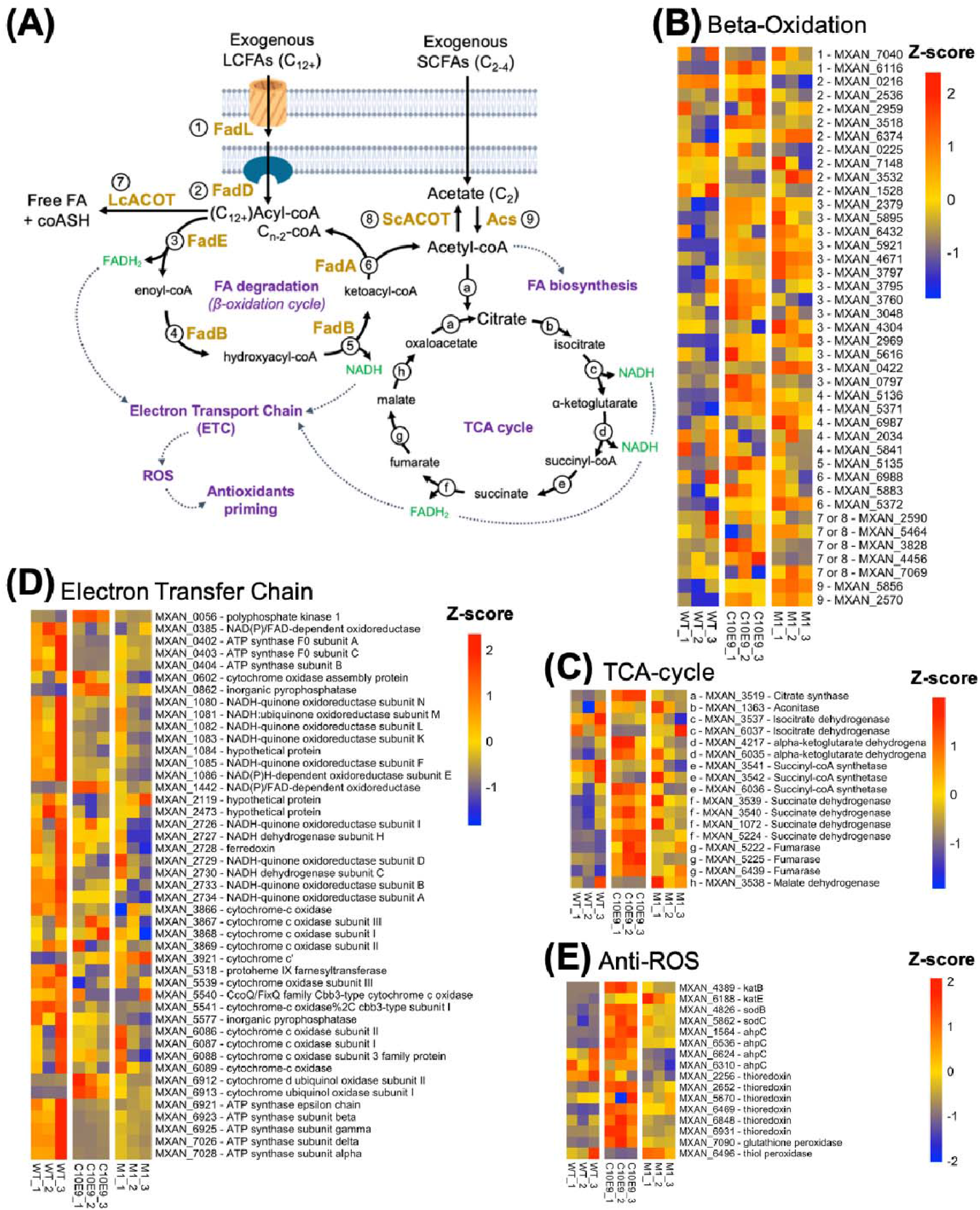
Metabolic Adaptation Mechanisms in Predator Mutants. **(A)** Predicted Exogenous FA-degradation pathway based on *M. xanthus* genome annotation. Long-Chain Fatty Acids (LCFAs, ≥ C12) are imported via the FadL transporter (1), while Short-Chain Fatty Acids (SCFAs, e.g., Acetate) diffuse directly across the membrane. LCFAs are activated by acyl-CoA synthase FadD (2) and subsequently undergo β-oxidation through enzymes FadE (Acyl-CoA dehydrogenase, 3), FadB (Enoyl-CoA hydratase and Hydroxyacyl-CoA dehydrogenase, 4 and 5), and FadA (Ketoacyl-CoA thiolase, 6), producing NADH and FADH2. In contrast, Acetate is directly activated to Acetyl-CoA by Acetate-coA synthetase (9). Longchain Acyl-CoA and Acetyl-CoA are deactivated by Long chain Acyl Thioesterase LcACOT and Short chain acyl-CoA ScACOT respectively (7 and 8). Acetyl-CoA enters both fatty acid biosynthesis and the TCA cycle, generating NADH and FADH2 for the Electron Transfer Chain (ETC). Depending on ETC activity, electron overflow can occur, leading to ROS production. High ROS levels initiate oxidative stress pathways, enhancing antioxidant production and modulating iron homeostasis. **(B-E)** Heatmap representations of gene expression profiles (RNA-seq) of genes annotated to be involved in Fatty acid β-oxidation (B), TCA cycle (C); Electron Transfer Chain (D) and Antioxidant defense (E). Numbers (1–9) preceding the gene MXAN accession numbers in (B) indicate their functional annotation with respect to the cycle shown in (A). Strains compared: *M. xanthus* DZ2 WT, C10E9, and M1. Each column within the heatmaps represents an independent biological replicate. The values shown in the heatmaps represent DESeq2normalized counts that have been scaled across conditions for each gene. This scaling allows for the visualization of relative expression level changes for multiple genes simultaneously. However, it’s important to note that this scaling does not accurately represent the absolute fold change differences between conditions.

Collectively, these results suggest that the evolution of super predator phenotypes is linked to the constitutive activation of a predatory metabolic program, possibly via a constitutive activation of the fatty acids β-oxidation pathway. Strikingly, the metabolic upshift observed in the C10E9 strain is mitigated in the M1 strain, suggesting that metabolic fine-tuning was key to the emergence of this phenotype (see Discussion).

### MXAN_5856 is an Acetyl-CoA Ligase and MXAN_2590 is an Acyl-CoA Thioesterase

The mutations in a putative Acyl-CoA Ligase and Acyl-CoA Thioesterase, two enzymes that act at the crossroad of fatty acids metabolism and the Krebs cycle (Figure 5A), further suggests the importance of metabolic changes driving enhanced predation. To demonstrate this, we explored the biochemical functions of each of these putative enzymes and tested whether their activity is indeed linked to Acetyl-CoA substrates as suggested by gene annotations. We expressed and purified MXAN_5856 (AcoL) and MXAN_2590 (AcoT), along with their respective W396* and M61I variants.

The MXAN_5856 protein is a predicted Acetate–CoA Ligase belonging to the AMP-forming Acetyl-CoA Synthetase family (ACS), which typically catalyzes the formation of Acetyl-CoA from Acetate and CoenzymeA precursors using ATP(24). We thus tested the activity of MXAN_5856 by monitoring ATP hydrolysis rates in the presence of free CoenzymeA (CoASH) and varying concentrations of Potassium Acetate (Figure 6A)(24). The purified AcoL protein showed high affinity for Acetate substrate (Km = 9,45 mM) with a catalytic efficiency (k_cat_/Km) of 2.64 nM^-1^.s^-1^, which is within the same range as the *Salmonella* AcsA enzyme (Km = 41 mM; k_cat_/Km = 0.23 nM^-1^.s^-1^)(25). In contrast, the catalytic efficiency of the W396* form (the M1 ACOL) was instead 0.07 nM^-1^.s^-1^, 27-fold lower than the WT enzyme (Figure 6A). This reduction was due to a 20-fold increase in Km, indicating that the W396* profoundly affects substrate binding and renders the enzyme mostly inactive in the evolved C10E9 and M1 strains (Figure 6A).

**FIGURE 6:**
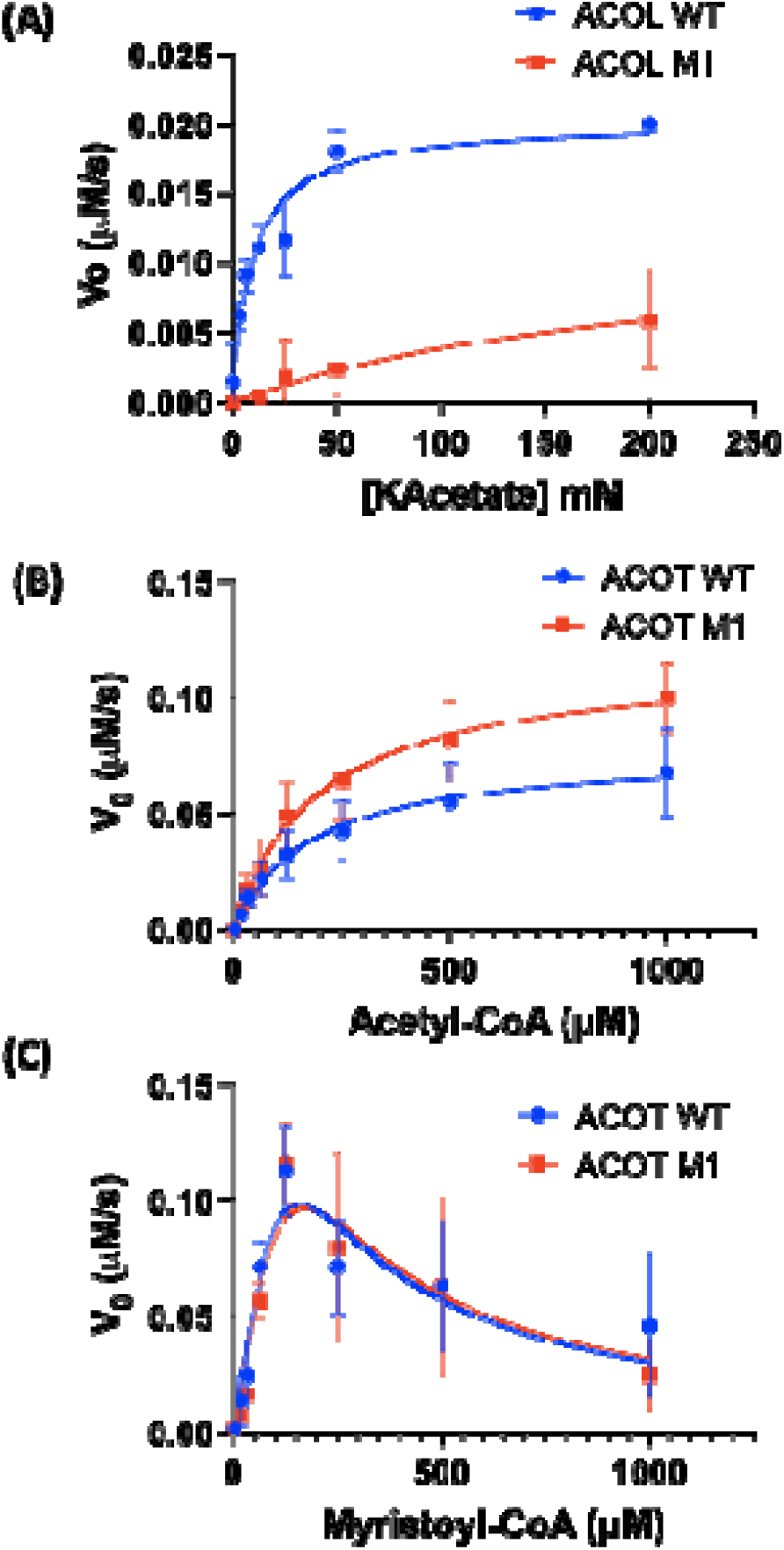
Enzymatic activities of ACOL and ACOT enzymes. **(A)** AMP-forming Acetyl-CoA Ligase activity of purified MXAN_5856 ACOL enzyme with Potassium Acetate (KAcetate) as varying substrate, and CoASH, ATP as fixed substrates; Michaelis-Menten representations are displayed; values are means ± S.D. of three replicates. **(B)** Acetyl-CoA Thioesterase activity of MXAN_2590 ACOT enzyme with Acetyl-CoA substrate. Values are means ± S.D. of four replicates. **(C)** Long chain Acyl-CoA Thioesterase activity of MXAN_2590 ACOT enzyme with Myristoyl-CoA substrate. Values are means ± S.D. of four replicates.

Next, we tested the biochemical activity of AcoT MXAN_2590. Hot-Dog fold Acyl-CoA Thioesterases hydrolyze fatty acyl-CoA intermediates, freeing CoASH from the acyl chains. In general, these enzymes control the intercellular balance between fatty acyl-CoAs and free fatty acids(26). To confirm that MXAN_2590 is an AcoT, we first tested its activity against Acetyl-CoA using the Ellman assay, following the formation of free CoASH(27). We observed that MXAN_2590 is a *bona fide* AcoT displaying a substrate affinity Km value for Acetyl-CoA at 180 μM (Figure 6B), which is in the same range as human ACOT12 (Km = 360 μM)(27). We also found that the point mutation M61I increases AcoT maximum velocity V_max_ by 52% (from 0,077 μM.s-1 to 0,117 μM.s-1), consistent with a gain of function (Figure 6B). We then tested its activity against long-chain acyl-CoAs using Myristoyl-CoA (C14- CoA) as a substrate. AcoT MXAN_2590 hydrolyzed C14-CoA, but the activity did not follow typical Michaelis-Menten kinetics and was inhibited at high substrate concentrations (Figure 6C), a typical behavior of substrate inhibition by micelles of long-chain fatty acyl-CoA on ACOT enzymes (28). On Myristoyl-CoA, the M61I point mutation did not provide a gain of activity, suggesting that *in vivo*, this mutation likely provides an advantage for the degradation of Acetyl-CoA and short-chain fatty acids (Figure 6C).

### Increased resistance to oxidative stress is key to the enhanced predatory performance of the M1 strain

In addition to fatty acids metabolism, RNA-seq also revealed that the expression of ROS-scavenging enzymes is highly upregulated, with extreme levels in the C10E9 strain, and more moderate but still high levels in the M1 strain compared to the WT (Figure 5E, quantified in Figure S6). In contrast, this oxidative stress response only becomes activated in the WT strain in the presence of prey, independently of the considered species (Figure S5B). This suggests that the constitutive expression of redox enzymes in the M1 strain protects it from ROS encountered during predation.

In the M1 strain, the catalase *katB* gene is expressed 50-fold higher than in WT (Figure S6, Table S3). The predicted KatB protein contains a signal sequence suggesting that it is secreted and protects against exogenous, rather than intracellular, hydrogen peroxide H_2_O_2_. To test this, we introduced exogenous catalase to the medium and analyzed the relative predatory performance of the WT and M1 strains. In the presence of catalase, the WT became just as efficient a predator as the M1, implying that heightened resistance to H_2_O_2_, one of the three primary ROS, plays a role in enhanced predatory performance (Figure 7A). This effect was not observed when catalase was heat-inactivated demonstrating that catalase activity relieves an oxidative stress burden on the predator (Figure 7A). Supporting this, WT and M1 strains showed similar growth kinetics in the presence of exogenous catalase (Figure S7).

**FIGURE 7:**
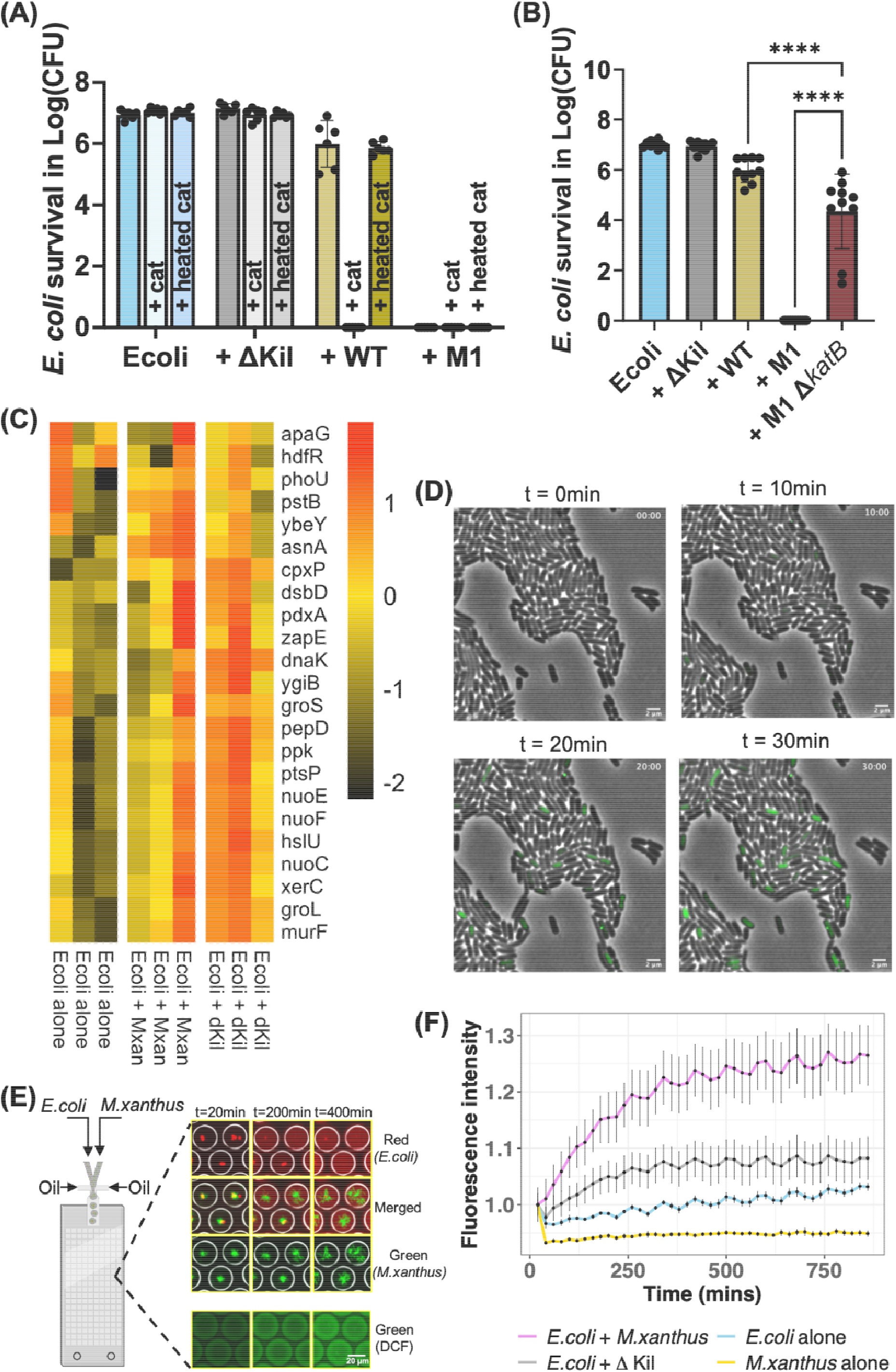
ROS adaptation explains the M1 phenotype. **(A)** Influence of catalase addition on the predation of *E. coli* MG1655 by *M. xanthus* DZ2 strains. The bar graph illustrates the mean survival of E. coli MG1655 in log10(CFU) after a 24-hour coincubation with M. xanthus DZ2 WT, predation-deficient ΔKil, and evolved mutant M1 predators, in the presence or absence of exogenously added, active or heat-inactivated catalase (5µg/ml). Each bar denotes the mean +/- SD from 3 biological replicates. Pairwise statistical comparisons were made using Mann-Whitney tests, with *p*-values indicated on the graph. **(B)** Deletion of *katB* in M1 diminishes its predation efficiency. The bar graph presents the average survival of *E. coli* MG1655, expressed in log10(CFU), after a 24-hour coincubation with various *M. xanthus* DZ2 predators: WT, predation-deficient ΔKil, evolved mutant M1, and a *katB*-deleted variant of M1. Each bar represents mean +/- SD derived from 3 biological replicates. Statistical significance was determined using one-way ANOVA, with subsequent multiple comparisons conducted using the Dunnett test; **** *p*< 0.0001. **(C)** Expression patterns (RNA-seq) of genes associated with hydroxyl radical production in *E. coli* subject to antibiotics stress(28) in response to WT *M. xanthus* and the Δ*kil* mutant strain of *M. xanthus*. Each column within the heatmaps represents an independent biological replicate. The values shown in the heatmaps represent DESeq2normalized counts that have been scaled across conditions for each gene. **(D)** Time-lapse fluorescence microscopy images revealing *E. coli* intracellular ROS accumulation upon interaction with the *M. xanthus* DZ2 ΔT3SS mutant, a strain that kills without lysing the prey. Before the interaction with the *M. xanthus* predator, *E. coli* prey cells had been incubated with the ROS-sensitive dye CM-H2DCFDA, which emits green fluorescence upon oxidation. Time points post-interaction are represented at 0, 10, 20, and 30 minutes. The displayed images are representative of two independent experiments. **(E)** Schematic of the droplet assay used to quantify oxidative stress produced by *E. coli* during predation. Suspensions of *E. coli* and *M. xanthus*, along with HFE7500 oil, were introduced into a custom droplet chip. Top: *Myxococcus* cells expressed GFP, and *E. coli* expressed mCherry. Water-in-oil microdroplets were generated using CF media. Over time, fluorescent *E. coli* cells disappeared while *Myxococcus* cells multiplied, indicating predation within the microdroplets. Bottom: In the oxidative stress experiment, *E. coli* was pre-incubated with CM-H2DCFDA, and neither *M. xanthus* nor *E. coli* were fluorescently labeled, allowing only the green fluorescence from oxidized CM-H2DCFDA to be recorded. Note that overtime, the droplets turn green as CM-H2DCFDA is released and oxidized **(F)** Quantification of CM- H2DCFDA oxidation in the droplets.The graph depicts mean intensity data derived from approximately 700 droplets per experiment, gathered across three distinct experiments, each spanning a 15-hour recording period. Fluorescent intensity values for each condition were normalized using the mean fluorescent value at the first time point to illustrate the relative increase in fluorescence over time. Each data point on the graph represents the average of three experiments, each encompassing around 700 droplets, with error bars indicating the standard error across these three experiments.

To directly demonstrate the importance of KatB, we introduced a Δ*katB* deletion in the M1 background. As expected, this mutation diminished predatory efficiency (Figure 7B). However, even with this reduction, the M1 Δ*katB* mutant still outperformed the WT, indicating that while *katB* contributes to enhanced predation, it is not the sole factor. This may not be surprising, given that other ROS detoxifying enzymes are also overexpressed in the M1 strain (Figure 5E, Figure S6). Interestingly, deletion of *katB* in the WT strain had no effect on predation, suggesting that it only becomes important in the evolved M1 background (Figure S8). Taken together, these observations confirm that ROS present a significant challenge for the predator during predator-prey interactions. Enhanced resistance to ROS, at least partially via KatB, emerges as an important factor contributing to the enhanced predation observed in the M1 strain.

### Predatory cells are exposed to the liberation of ROS upon contact-dependent lysis

What is the source of the ROS experienced during predation? Bacteria that are exposed to lethal attacks, antibiotics but also Type-6 secretion-mediated contact-dependent killing tend to produce ROS(28–30). If this also occurs when *Myxococcus* attacks *E. coli*, a ROS outburst could happen upon prey cell lysis. To test this, we conducted an RNA-seq experiment to monitor gene expression in *E. coli* exposed to *M. xanthus* and measured the expression of genes typically induced by the stresses mentioned above(28, 30). Incubating *E. coli* with *M. xanthus* WT induced the expression of genes that are also activated in response to antibiotics (Figure 7C)(28). However, this response was variable, which we reasoned might be due to the difficulty of measuring gene expression robustly in actively predated cells. We therefore performed a similar experiment using a *Myxococcus* Δ*kil* mutant that can invade *E. coli* colonies but cannot kill *E. coli* cells by contact. The *M. xanthus* Δ*kil* mutant induced a robust response, suggesting that interactions between *Myxococcus* and *E. coli* lead to general stress and ROS manifestation (Figure 7C). Interestingly, this response is not Kil-dependent.

Next, we aimed to image ROS production in *E. coli* cells directly when they are in contact with *Myxococcus* cells. For this, we used *E. coli* cells pre-labeled with CM-H2DCFDA, a general ROS probe that emits green fluorescence upon oxidation (Figure S9)(29, 31). As expected, the pre-labeled *E. coli* were non-fluorescent in the absence of predator contacts. However, following contact, the *E. coli* cells lysed within minutes, making it difficult to monitor ROS accumulation. Occasionally, a sporadic fluorescence surge was noticeable in *E. coli* cells adjacent to the lytic contact (Figure S10), suggesting that cell rupture may discharge a ROS wave impacting neighboring cells. However, this was rarely observed, possibly due to the swift diffusion and dilution of ROS in the extracellular environment.

To definitively confirm that cell-cell contacts cause an oxidative surge, we adopted two complementary strategies. First, we used a *Myxococcus* strain with a functional Kil system but lacking the T3SS (Δt3ss). Cells carrying this genotype establish prolonged contact with the prey cell, inducing plasmolysis but without causing cell rupture (12). Thus, if a ROS build-up occurs in the prey cells, they should become fluorescent when exposed to the Δ*t3ss* mutant. As shown in Figure 7D, *E. coli* cells preloaded with CM-H2DCFDA and initially non-fluorescent turned bright following contact with *Myxococcus* cells, indicating a ROS burst in prey cells.

Since this effect could be argued to be specific to the Δ*t3ss* mutant, we designed a second strategy to visualize ROS liberated following *E. coli* lysis by the WT strain. We engineered a novel predator-prey interaction assay within microdroplets. This setup provided a confined micron-sized compartment, limiting the dilution of CM-H2DCFDA when released upon prey cell lysis. Thus, if ROS are produced in the droplets, fluorescence should accumulate. Killing of *E. coli* cells and growth of *Myxococcus* cells were observed within the microdroplets, albeit at a slower pace (Figure 7E). When droplets were loaded with CM-H2DCFDA-labeled *E. coli* alone, only weak fluorescence was observed, consistent with low ROS levels in the absence of the predator. However, when CM-H2DCFDA-labeled *E. coli* cells were co-incubated with *Myxococcus* WT strain, a significant, time-dependent fluorescence increase was detected within the droplet (Figure 7E, 7F). Consistent with the *E. coli* RNA-seq analysis, we also observed fluorescence increase with the Δ*kil* mutant, though at a lower level than with WT, likely because contact-dependent killing liberates ROS that build up in the prey cells in the droplets.

Therefore, Kil-dependent killing produces an oxidative burst in prey cells, which may impose stress on predator cells upon cell lysis. This could, at least in part, explain the superior performance of the M1 variant, where predator cells are already primed with ROS-scavenger production.

## Discussion

Despite growing evidence that bacterial predators are ubiquitous in the environment, little is known about the mechanisms by which they interact with and consume their prey. Evolution experiments have the potential to reveal critical features of predator-prey interactions, mechanisms of prey resistance, and specific genes involved in these interactions(32–35). In a seminal study, Nair and Velicer conducted a co-evolution experiment where both predator and prey (*E. coli*) were observed to co-adapt via accelerated genomic evolution specific to each species(35). Remarkably, they observed the parallel evolution of predator (*eatB*) and prey (*ompT*) loci, suggesting that these genetic traits are important in predator-prey interaction. In this study, we did not allow the prey to evolve and instead focused on the selection of enhanced predatory phenotypes. This approach was uniquely powerful in identifying new genetics determinants of predation and revealing important aspects of the metabolic interactions.

First, we discovered that augmented predatory phenotypes result from profound changes in the expression of the genetic repertoire. It is striking that both the C10E9 and M1 strains constitutively activate genes that are induced by the presence of prey in the WT. Such a large genetic shift implies the existence of a prey-specific program, which could occur via the modification of a central regulator. We hypothesize that the gain-of-function mutation in Histidine Kinase *EpsJ* is at least partially responsible for this genetic switch. Consistent with this, in our evolution experiment, we identified five other enhanced predators distinct from the M1 strain (to be presented in a follow up study), all carrying a D79N substitution in EpsJ. This strongly suggests that this mutation acts as a bottleneck for the evolution of predatory phenotypes. Thus, defining the EpsJ regulon and elucidating the function of downstream genes opens exciting future perspectives for understanding the induction and structure of the genetic predation program, and is a focus of our ongoing research.

We stress however that the *epsJ* mutation alone is not sufficient for the emergence of a selectable phenotype, highlighting the importance of combinatorial effects in the evolution of predatory phenotypes. We determined, genetically and biochemically, that enzymes involved in (i) remediating oxidative stress and (ii) regulating acetyl-CoA fluxes and fatty acid metabolism are critical for heightened predatory performance. Remarkably, the oxidative stress response and the fatty acid β-oxidation pathway are among the most highly induced genes when the evolved strains are compared to the ancestor. These genes are also induced when the ancestor is mixed with prey, independent of the prey species. Thus, we propose that changes in fatty acid metabolism and oxidative stress response are two major facets of predator evolution.

*Predator metabolism*. The precise mechanism by which loss of function in *AcoL*, a *bona fide* AcsA-like enzyme responsible for producing acetyl-CoA from acetate and CoA, enhances predation remains unclear. Notably, *AcoT*, an enzyme that catalyzes the reverse reaction, is essential for predation. Interestingly, the M61I mutation in *AcoT* appears to further deplete acetyl-CoA pools through a gain-of-function effect that increases acetyl-CoA hydrolysis. This effect is not observed with long-chain fatty acid intermediates, such as myristoyl-CoA. The beneficial effect of the M61I mutation may therefore not involve diverting long-chain fatty acids away from the β-oxidation cycle for alternative processes, such as membrane synthesis. Instead, it may directly impact acetyl-CoA pools. Acetyl-CoA is a central metabolite at the crossroads of multiple metabolic pathways, including the TCA cycle. By reducing acetyl-CoA levels, these mutations might help prevent metabolic overload and allow the cell to more effectively process prey-derived nutrients. Indeed, these mutations are clearly linked to metabolic changes: for instance, while TCA cycle gene expression is upregulated in the C10E9 strain, it is downregulated in the M1 strain, suggesting that the M61I mutation mitigates this effect. This could be even more important when the predator is utilizing lipids from the prey, as fatty acid β-oxidation generates acetyl-CoA. Lowering acetyl-CoA levels might facilitate the continuous flow of β-oxidation by preventing product inhibition of 3- oxoacyl-CoA thiolase, catalyzing the final thiolytic step of the cycle(36). The exact *in vivo* consequences of the mutations will require comprehensive metabolic analyses, because the *M. xanthus* genome encodes additional predicted *AcoL* and *AcoT* enzymes, which could introduce counterbalancing effects.

*Oxidative stress*. Several lines of evidence demonstrate that ROS emitted by the prey cells upon contact with the predator generate an important burden for the predator:

i. Addition of extracellular catalase cancels the advantage of the M1 strain, demonstrating that its performance is mostly driven by its resistance to oxidative stress in our conditions
ii. A fluorescent ROS indicator shows that ROS build up in prey cells upon contact and are released into the extracellular environment upon prey lysis. Consistent with ROS as an exogenous stress, KatB, is predicted to be secreted (based on its signal peptide), and it serves as a major protective enzyme in the M1 strain. In *P. aeruginosa*, a similar extracellular catalase (KatA) provides resistance to oxidative stress during infection(37).
iii. ROS are likely generated as part of a stress response by the prey cell in the presence of the predator. This stress response might be similar to what has been described when *E. coli* is attacked by the Type-6 secretion system (T6SS), another mode of contact-dependent killing(29). While the exact pathway leading to ROS production in prey remains to be identified, analysis of prey transcription reveals that genes commonly induced after antibiotic stress are also induced upon predator contact, suggesting a common stress signature.

In conclusion, the metabolic changes observed in the evolved strains echo emerging research suggesting that the fatty acid degradation pathway might enhance bacterial stress responses, better equipping them to navigate challenging environments(38). Predator-prey cell interactions involve both fatty acid metabolism and oxidative stress defense, drawing interesting parallels to pathogen-host cell interactions.(39, 40). However, we have no evidence here that ROS production is a defense mechanism of the prey; it more likely represents a stress response, as discussed above. A limitation of this study is that it was conducted in a laboratory setting, which may limit the ecological relevance. The M1 genotype might carry additional constraints that could prevent the emergence of this variant in more natural habitats. Alternatively, the wild-type strain may have adapted to prey-independent growth through domestication and prolonged cultivation in nutrient-rich media. Nevertheless, our findings suggest that a spectrum of predatory phenotypes can emerge in response to specific environmental conditions. In the future, sequencing wild strains directly sampled from soil environments could provide valuable insights into the occurrence of mutations similar to those observed in the M1 strain.

## Acknowledgements

We would like to thank Benjamin Ezraty for helpful discussions; Emma Bouveret for helpful discussions and help with fatty acids metabolism gene annotation; Julien Herrou for initial discussions at the beginning of the project and help with experiments; Swapnesh Panigrahi for helping with image analysis. Guilhem Chenon for assistance in setting up the droplet-based assays; Beatrice Py, Corinne Aubert and Mathieu Sourice for fruitful discussions. Sophie Helaine for helpful comments on the manuscript.

## Author contributions

Conceived and designed the experiments: R.J., N.-H.L., B.H. and T.M.. Performed the experiments: R.J., N.-H.L., L.B., J.Ba. and J.Bi.. Analyzed the data: R.J., N.-H.L., L.B., Y.D., B.H., and T.M.. Wrote the paper: R.J., N.-H.L., and T.M. All authors commented on the manuscript. **Competing interests:** All other authors declare that they have no competing interests.

## Data and materials availability

All data needed to evaluate the conclusions in the paper are present in the paper and/or the Supplementary Materials.

## METHODS

### Bacterial strains and growth conditions

*M. xanthus* DZ2 strains were cultivated in Casitone Yeast Extract (CYE) medium, while *E. coli* MG1655 and *B. subtilis* strain 168 cells were grown in Luria-Bertani (LB) broth. The *C. crescentu*s strain NA1000 was cultured in PYE (Peptone Yeast Extract). All of the strains were grown at 32°C and the experiments were performed on 1.5% hard agar plates.

### Effect of glucose concentration on predation

*M. xanthus* and *E. coli* were cultured overnight in 20 mL of their respective growth media at 32°C. On the following day, the cells were collected by centrifugation and resuspended in CF medium (comprising MOPS 10 mM at pH 7.6, KH2PO4 1 mM, MgSO4 8 mM, (NH4)2SO4 0.02%, Na citrate 0.2%, and Bacto Casitone 0.015%). The OD600 of both cultures was adjusted to 5. Subsequently, 10 µL of the *E. coli* suspension was carefully deposited alongside 10 µL of the *M. xanthus* suspension onto separate CF agar plates, each with varying glucose concentrations (0%, 0.075%, 0.1%, and 0.2%). Once the drops had air-dried, the plates were incubated at 32°C, and observations were made after a 72-hour incubation period (Figure 2A).

### Evolution experiment

*E. coli* was grown overnight in 20 mL of LB at 32°C. The following day, the cells were pelleted and resuspended in CF medium with the OD600 adjusted to 5. 20 µL of this suspension was then placed on a CF 1.5% agar plate containing 0.075% glucose. After the drop had dried, *M. xanthus* cells (previously grown on CYE agar plates at 32°C) were added to the center of the *E. coli* drop using a sterilized pipette tip. After about 6 days, when the *M. xanthus* cells had consumed all of the *E. coli* and reached the edge of the *E. coli* colony, they were removed from the edge of the colony and transferred to a new 20 µL *E. coli* drop (Figure 2B). This cycle was repeated 20 times, and after every 5 cycles, the concentration of glucose in the CF agar was increased by 10% of the previous concentration. Additionally, after each 10 cycles, a DMSO stock of the evolved *M. xanthus* was prepared.

### Construction of *M. xanthus* mutant strains

All primers, plasmids and strains used for this study are listed in Tables S5-S7. Gene deletions and point mutations in *M. xanthus* were achieved using a double-recombination approach as described previously(41). In brief, sequences ranging from 800 to 1000 nucleotides flanking the targeted genes for deletion (or the point mutations for insertion) were PCR amplified and assembled into the pBJ114 suicide plasmid (GalS, KanR) via Gibson assembly. This plasmid was then electroporated into *M. xanthus*. Following successive selections with Kanamycin and Galactose, clones harboring the desired genetic modifications were identified using PCR.

### Predation killing assay

*Myxococcus* and kanamycin-resistant strains of the prey (*E. coli*, *B. subtilis*, or *C. crescentu*s) were grown overnight in 20 mL of their respective rich media at 32C. Cells were pelleted and the OD600 of *Myxococcus* and prey was adjusted to 1 and 5, respectively, in CF medium. The *Myxococcus* and prey were mixed in a 1:3 ratio, and a 10 µL drop of this mixture was placed on a CF 1.5% agar plate containing 0.075% glucose. After 24 hours, cells from the spot were harvested and the colony forming units of prey were counted on Kanamycin-supplemented LB media for *E. coli* and *B. subtilis*, and on PYE supplemented with Kanamycin media for *C. crescentus*. Each assay is performed in technical duplicate.

### Growth Curve Determination of *M. xanthus* via Digital PCR (dPCR)

To assess the growth of *M. xanthus* during *E. coli* predation on CF agar media, we employed droplet digital PCR (dPCR) to directly quantify its genomic DNA concentration, using it as a surrogate measure for growth. Genomic DNA was extracted from three pooled killing assay spots (as detailed in the previous section) utilizing the Monarch® Genomic DNA Purification Kit (NEB). Quantification of genomic DNA copies was targeted at the *kdpAB* gene and carried out with the Naica® Crystal Digital PCR System using Sapphire chips (Stilla Technologies).

The dPCR reactions consisted of PerfeCta® qPCR ToughMix® UNG (Quantabio), 0.8 ng/µL Alexa 647 (Invitrogen), 1.5X EvaGreen® (Biotium), 4% DMSO (Biolabs), 200 nM of each primer *kdpAB*(37), and 10 µL of the DNA template. Amplification was performed on the Geode thermocycler (Stilla Technologies) under the following conditions: an initial 95°C for 3 minutes, succeeded by 45 cycles of 95°C for 10 seconds and 60°C for 15 seconds. Fluorescence acquisition was accomplished using the Prism3 imager and Crystal Reader software (Stilla Technologies), with data analysis conducted via the Crystal Miner software (Stilla Technologies).

### Protein production and purification

For expression and purification of AcoL (MXAN_5856), AcoT (MXAN_2590) and their mutant variants, the corresponding genes were amplified from either *M. xanthus* WT or M1 strain. The amplicons were inserted in the pET28a+ vector (Novagen) at the NdeI and HindIII restriction sites. *E. coli* LEMO21 producing strain harboring the constructed vectors was used to produce the N-terminal His-tagged proteins in auto-inducible media ZYM-2025 for 48h at 20°C. Cell pellet were resuspended in Lysis Buffer containing 50mM Tris-HCl pH 8.0, NaCl 300mM, Imidazole 20mM, MgCl2 10mM, DNAse 10mg/ml and Lysozyme 10mg/mL, then lysed using Emulsiflex-C5 (Avestin). The lysates were clarified by centrifugation at 10,000 x g for 15 min. Clarified lysates were incubated with Ni-NTA beads and the recombinant proteins were eluted at 180mM Imidazole. Eluted fractions were desalted using PD-10 columns (Merck) then concentrated to around 15mg/ml. Purified proteins were stored in 50% Glycerol at -20°C.

### Acetate-CoA Ligase AcoL enzymatic assay

The AMP-forming AcoL (MXAN_5856) WT and mutant variant activities were measured by quantifying the pyrophosphate co-product in a pyrophosphatase coupled assay. Assays were performed in 96-well microplates (Greiner Bio-One) in 100 μl of reaction mix containing 100mM Tris-HCl (pH 8.0), 4mM MgCl2, 2.5mM ATP, 0.5mM 1,4-dithiothreitol, 2 milliunits/ml pyrophosphatase (Sigma), and varying concentration of Potassium Acetate (Sigma). Reactions were started by adding 800 nM AcoL in the assay and incubated 30 min at 32°C. A reaction without substrate was performed in each experiment and used as blank. The released phosphate products were quantified using the PiColorLock Gold photometric kit (Novus Biologicals) as instructed.

For apparent kinetic parameter determinations (Km, V_max_, and k_cat_), the initial velocities were calculated using initial absorbance (at 615 nm) reading following protein addition and after 30 min incubation, using TECAN plate reader. The Michaelis-Menten curves were fitted by non-linear regression analysis using GraphPad Prism 10.

### Acetyl-CoA Thioesterase AcoT enzymatic assay

The AcoT (MXAN_2590) WT and mutant activities were measured by quantifying the free CoenzymeA (CoASH) product with Ellman’s assay(42). Assays were performed in 96-well microplates (Greiner Bio-One) in 100 μl of reaction mix containing 100mM Tris-HCl (pH 8.0), 0.5mM 5,5′-dithiobis-(2-nitrobenzoic acid) DTNB (Sigma), and varying concentration of Acetyl-CoA or Myristoyl-CoA (Roche). Reactions were started by adding 25 nM AcoT in the assay a. A reaction without substrate was performed in each experiment and used as blank. The assay absorbance was read at 412 nm every 2.5 min for 30 min at 25°C using TECAN Plate reader. The initial velocities were determined using linear regression of the linear portion of the enzymatic velocity. The Michaelis-Menten curves were fitted by non-linear regression analysis using GraphPad Prism 10.

### RNA-seq library preparation and data analysis

*M. xanthus* strains were grown in 20 mL of CYE, while *E. coli* and *B. subtilis* cells were cultured in 20 mL of LB. The *C. crescentu*s was grown in 20 mL of PYE. All of the strains were grown at 32°C overnight. The next day, the cells were pelleted and the growth media was removed. The cells were resuspended in CF medium such that the OD600 of *M. xanthus* was adjusted to 5, and the OD600 of the prey bacteria was adjusted to 10. *M. xanthus* and prey bacteria were mixed in a 1:3 ratio, and 20 µL of this mixture was spotted onto a CF 1.5% agar plate containing 0.075% glucose. For the *Myxococcus* alone condition, an *M. xanthus* culture with an OD600 of 5 was mixed with CF medium in a 1:3 ratio. The plate was incubated at 32°C. After ∼2 hours, cells from 25 of the spots were collected using sterile loops, combined to form a single sample, and rapidly frozen in liquid nitrogen for storage at - 80°C. To extract RNA, the cells were lysed using lysozyme (10 mg/mL) and then the RNA was extracted using the TRiZol (Invitrogen) protocol. The total RNA was treated with TURBO^TM^ DNase (Invitrogen), and the quantity and quality of the RNA were measured using a Qubit and a TapeStation, respectively. Libraries were prepared using 1 µg of total RNA according to the Zymo-Seq RiboFree Total RNA Library Kit protocol. The quantity and quality of the libraries were also measured using a Qubit and a TapeStation. The libraries were sequenced on an Illumina NextSeq500 platform at TGML in Marseille, generating approximately 10-35 million 75 bp paired-end reads per sample. The reads were mapped to the *M. xanthus* DK1622 genome (GCF_000012685.1, NCBI) OR *E. coli* str. K-12 substr. MG1655 (GCF_000005845.2, NCBI) using Burrows-Wheeler Aligner (version 0.7.10)(43) default parameters and the number of reads mapping to each coding sequence (CDS) was counted using HTseq (version 0.6.1)(44). The read counts were then normalized utilizing the DESeq2 method(45), and genes exhibiting differential expression were identified with a p-adjusted value <0.05 using the Wald test in DESeq2. Gene ontology enrichment analysis was performed on differentially expressed genes using *clusterprofiler* 4.0 package(46) in R. The normalized read counts from DESeq2 were further utilized to create heatmaps illustrating gene expression patterns (Figure 4). To accomplish this, the DESeq2 normalized read counts were individually scaled for each gene across various conditions using the scale function in R. These scaled values were plotted using the *pheatmap* function in R. The genes related to beta-oxidation, the TCA cycle, the Electron Transfer Chain, and ROS response, as shown in heatmaps, were annotated using a combination of the KEGG database, literature search, and a manual homology search to identify *E. coli* homologs within the *Myxococcus* genome.

### DNAseq library preparation and data analysis

*M. xanthus* strains were grown in 20 mL of CYE at 32°C overnight. The next day, 500 µL of the cultured cells were pelleted and the growth media was removed. DNA was extracted using the MasterPure DNA purification kit (Cat No. 85200) from Epicentre. NEBNext® Ultra™ II DNA Library Prep Kit for Illumina® (NEB #E7645S/L) was used for library preparation and libraries were sequenced on Illumina MiSeq or NextSeq 2000 platform and more than 1.25M read pairs of 150 bp length were generated per sample i.e. more than 40X coverage for *M. xanthus* DZ-2 genome. SNPs were detected using breseq version 0.35.1(47).

### Fluorescence Microscopy of ROS Accumulation in *E. coli* during *M. xanthus* Predation

*E. coli* prey and *M. xanthus* predator strains were cultured overnight in their designated media. The *E. coli* culture was subsequently diluted 100-fold in LB supplemented with 20 µM CM-H2DCFDA, an ROS-sensitive fluorescent dye, then grown to an OD600 of 2 at 37°C with shaking. After centrifugation, *E. coli* cells were resuspended in CF medium to an OD600 of 2. To test if dyed *E. coli* cells could reflect intracellular ROS level, 75 µL of cells were mixed together in black 96-wells plate (Greiner Bio-One) with 75 µL of either CF medium alone or CF medium containing 20 µM of redox-cycling compound phenazine-methosulfate (PMS). Fluorescence intensity of each well was monitored using the TECAN plate reader (Ex/Em = 485/520 nm).

Equal aliquots of *M. xanthus* and dyed *E. coli* suspensions were mixed and 1 µl was spotted onto a fresh CF 1% agarose pad on a microscope slide. Following a brief drying period and incubation in the dark at room temperature for 30 minutes, imaging was performed. The TE2000-E-PFS (Nikon) inverted epifluorescence microscope, equipped with a ×100/1.4 DLL objective and Nikon’s NIS software, was used. Time-lapse recordings of 30 minutes (capturing every 20 seconds) were conducted at x100 magnification. DIA images utilized a 20 ms light exposure, while GFP images required a 100 ms fluorescence exposure with a 30% power intensity at an excitation wavelength of 470 nm, ensuring minimized photobleaching and phototoxicity. The ’Perfect Focus System’ (PFS) was engaged throughout for consistent focus.

### Droplet microfluidic & Image analysis

*M. xanthus* and CM-H2DCFDA-dyed *E. coli* cells were prepared as described above. The dyed *E. coli* suspension was washed and adjusted to an OD600 of 2, while the *M. xanthus* suspension was set to an OD600 of 7. These distinct cell suspensions in CF media were introduced into separate sample inlets, along with another inlet for HFE7500 oil containing 2% weight/weight of FluoroSurfactant (RAN Biotechnologies, MA 01915), within a custom-designed droplet chip at ESPCI(48) (Figure 7E). Both CF and oil phases were drawn through orthogonal channels inside the chip, accomplished by generating a vacuum within the chip using a 5 mL syringe pump. This specialized chip facilitated the creation of thousands of uniformly sized droplets, each with a volume of around 80 pL, utilizing the introduced cell cultures. These droplets consisted of CF media suspended within an oil medium, establishing a water-in-oil droplet configuration. Each of these droplets contained a mixture of both *M. xanthus* and *E. coli* cells, with approximately 20 cells of each type present in every droplet. This cell composition was determined based on the previously specified OD values.

After the generation of droplets, they were pulled into a chamber with a consistent height of 30 µm, causing the droplets to undergo deformation into a flattened, pancake-like configuration. This compression facilitated their observation under a microscope. The compressed droplets exhibited an approximate diameter of 40 µm. These droplets were then subjected to imaging using an inverted epifluorescence microscope, specifically the TE2000- E-PFS by Nikon, operating at a 20x magnification. At this magnification level, each image frame could encompass approximately 700 droplets. A series of time-lapse recordings were performed, spanning a 15-hour duration, with images captured at 20-minute intervals throughout this period. To extract meaningful data from these images, a segmentation process was applied to delineate individual droplets, and subsequent analysis was carried out using ImageJ. This analysis aimed to calculate the mean intensity of each individual droplet.

### Statistical Analysis

Statistical analyses were performed using GraphPad Prism 10 (GraphPad Software Inc., La Jolla, CA) and R. Prism 10 was used to determine statistical significance. Non-parametric Mann-Whitney tests were performed to compare ranks between two conditions. One-way ANOVA analysis with post-hoc multiple comparison using the Dunnett test weas used on datasets with 3 or more conditions.

## Data Availability

The raw transcriptome sequencing data, in the form of raw reads, have been deposited in the NCBI Sequence Read Archive (SRA) with the Bioproject accession number PRJNA1040215.

---

**FIGURE S1:**
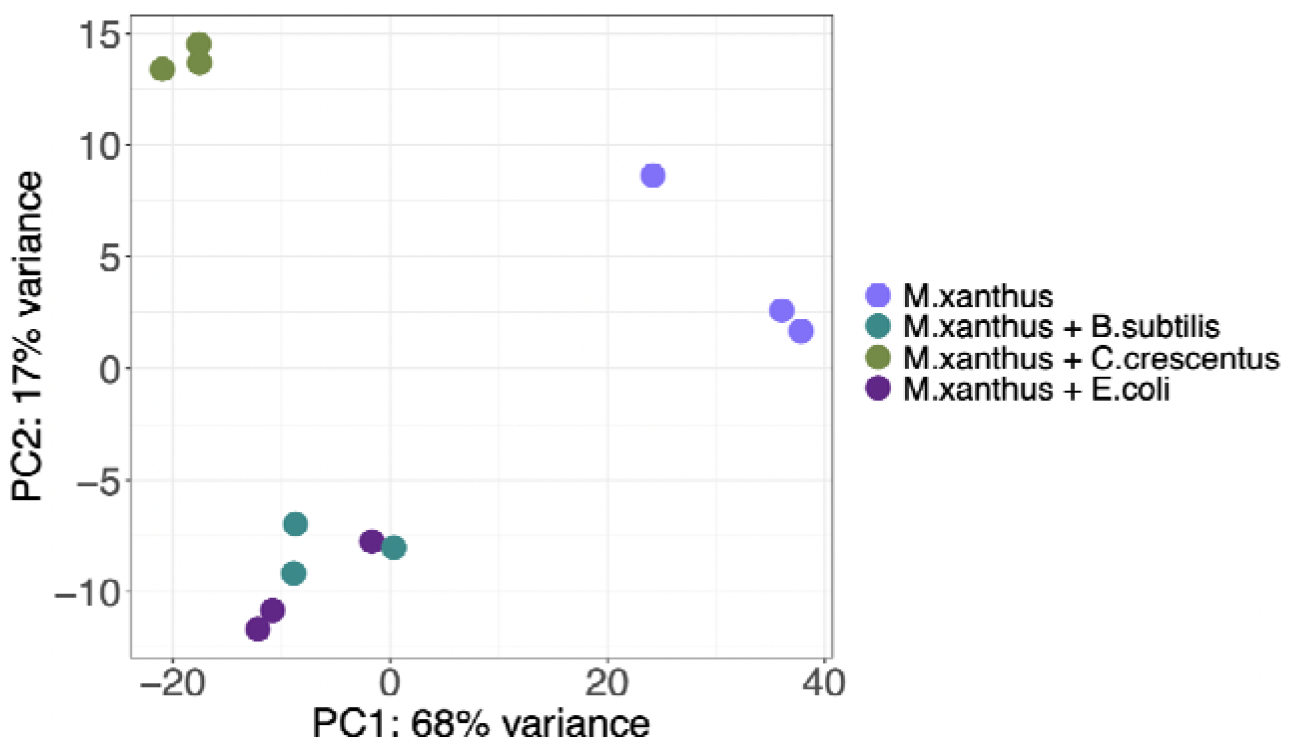
Transcriptomic response of *M. xanthus* WT strian to 3 different preys. Principal Component Analysis (PCA) was performed on all detected genes in RNA-seq, demonstrating the consistency among replicates. Notably, the transcriptomic response to *C. crescentus* differs significantly from that of *E. coli* and *B. subtilis*, while the responses to *E. coli* and *B. subtilis* appear highly similar.

**FIGURE S2:**
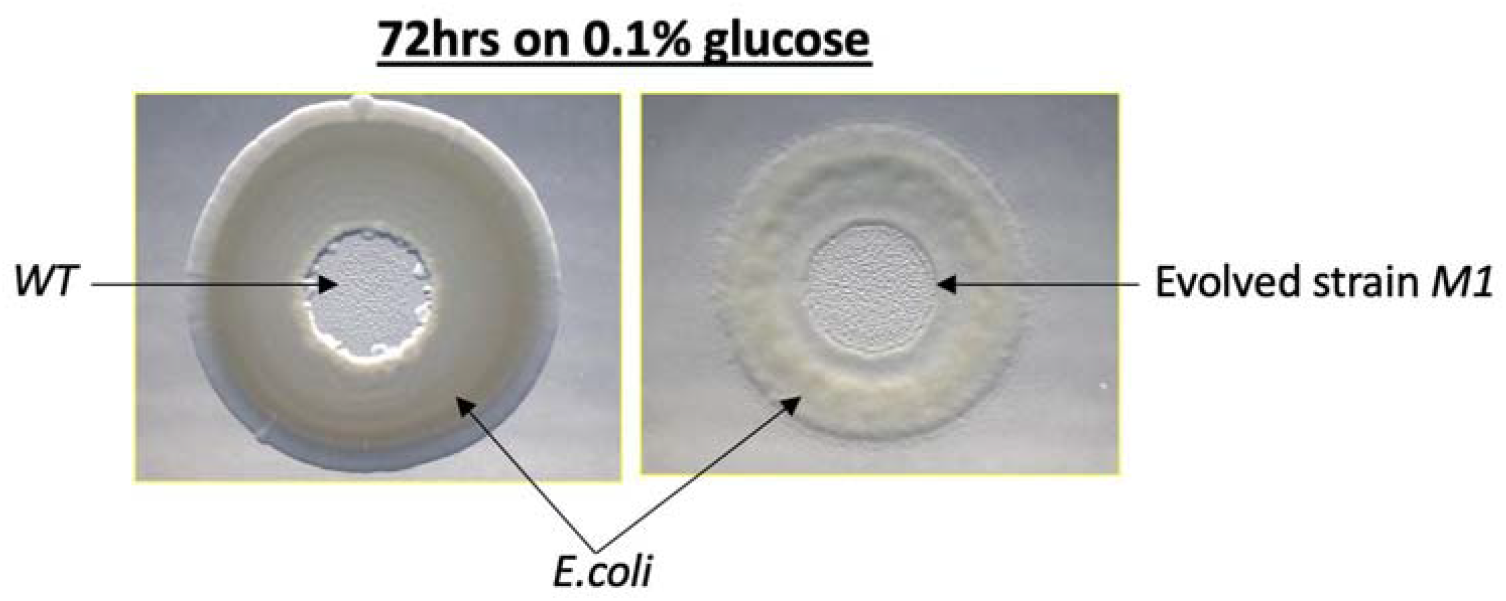
Evolved mutant M1, can kill prey on 0.1% glucose media. A 20 μL drop of *E. coli* cells, with an OD600 of 5, was deposited on 1.5% agar containing 0.1% glucose and allowed to air-dry. Subsequently, 1 μL (OD600 = 5) of *Myxococcus* cells (WT or M1) was carefully deposited at the center of the previously dried *E. coli* drop. Following a 72-hour incubation period, the M1 strain demonstrated impressive spreading capabilities, effectively colonizing and clearing the entire *E. coli* colony, whereas the WT strain failed to exhibit similar spreading behavior.

**FIGURE S3:**
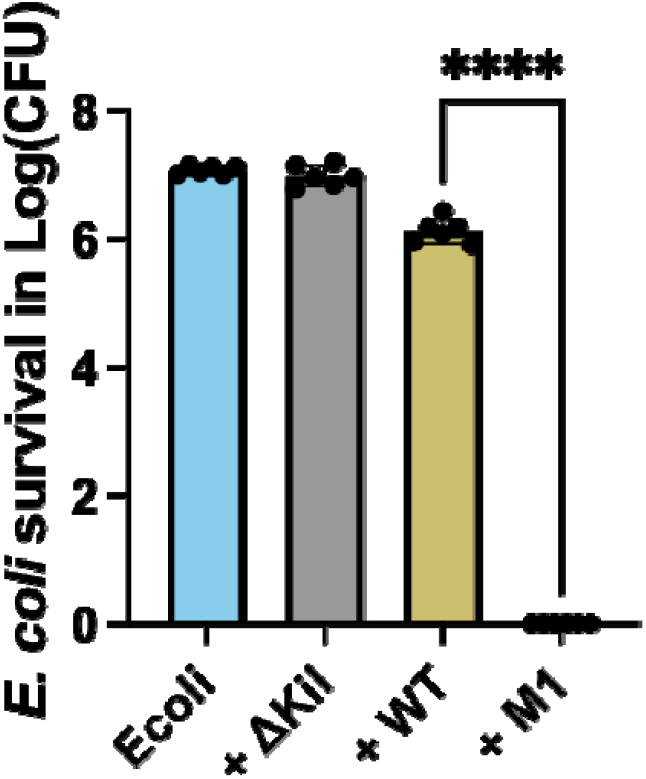
Predation efficiency of evolved mutant M1 compared to WT strain in buffered medium. The bar graph presents the average survival of *E. coli* MG1655, expressed in log10(CFU), after a 24-hour coincubation on CF media buffered at 100mM MOPS pH 7.5, with different *M. xanthus* DZ2 predators: WT, predation-deficient ΔKil, and M1 variant. Each bar represents mean +/- SD derived from 3 biological replicates. Statistical significance was determined using one-way ANOVA, with subsequent multiple comparisons conducted using the Bonferroni test; *\*\*\*\*p<0,0001*.

**FIGURE S4:**
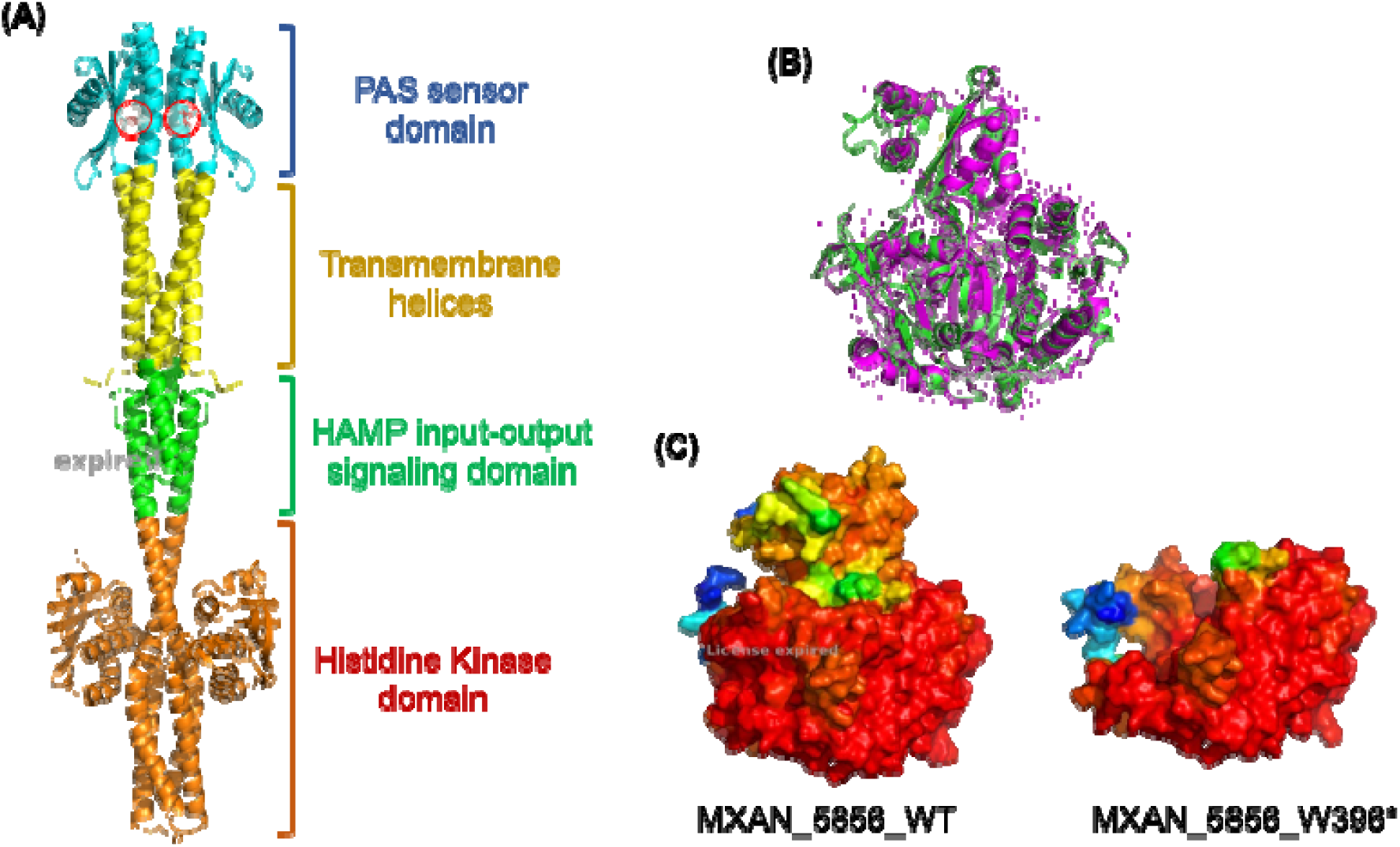
Structural Prediction of *M. xanthus* EpsJ (MXAN_7439) and ACOL (MXAN_5856). A) AlphaFold3-predicted model (pTM = 0.46) for *M. xanthus* EpsJ histidine kinase in a typical dimer formation. In cyan: Per-Arnt-Sim (PAS) sensor domain (residues 40–159). In yellow: transmembrane helices (residues 1–39; 160–191). In green: HAMP input-output signal domain (residues 192–244). In orange: histidine kinase domain (residues 245– 501). Red circles indicate D79 resides. (B) Superimposition of the AlphaFold3-predicted model (pTM = 0.96) for *M. xanthus* ACOL (MXAN_5856, green) with the experimentally resolved 3D structure of *S. enterica* AcsA (PDB: 1PG4, magenta). (C) AlphaFold3-predicted structures of the WT ACOL (left) and the M1 mutant variant (right, pTM = 0.96), showing the absence of the C-terminal domain in the M1 variant, which impairs enzyme activity.

**FIGURE S5:**
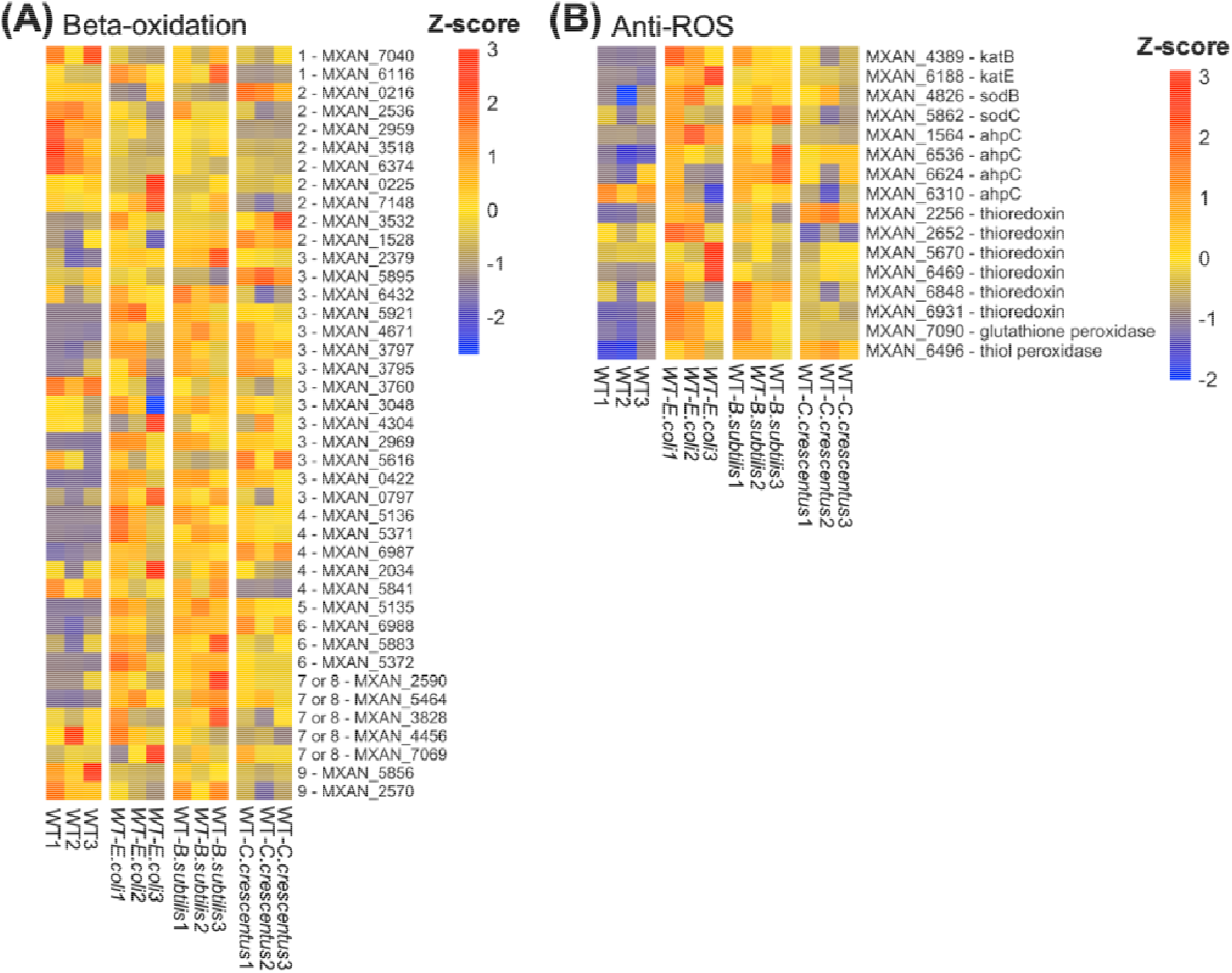
**Changes in gene expressions in *M. xanthus* WT during predation**. Heatmaps display gene expression profiles (RNA-seq), utilizing scaled DESeq2-normalized counts, with a focus on specific genes in *M. xanthus* as shown in Figure 4: **(A)** Genes related to fatty acid degradation (β-oxidation) **(B)** Antioxidant defense during predation. Conditions compared in (A) and (B) include *DZ2* WT alone, WT + *E. coli*, WT + *B. subtilis*, and WT + *C. crescentus*. Each column in the heatmaps represents an independent biological replicate.

**FIGURE S6:**
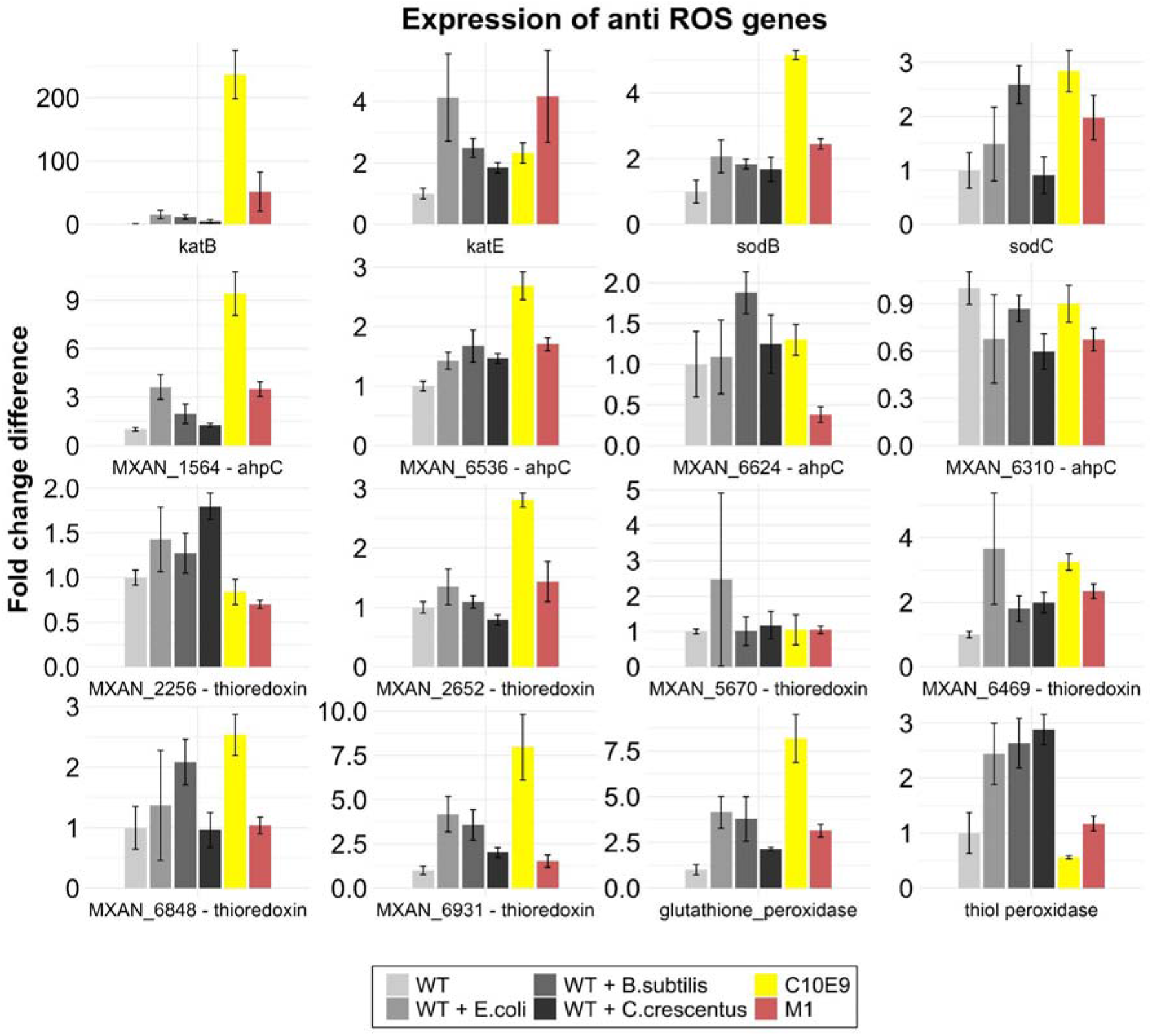
Antioxidant gene expression patterns during predation and evolved mutant variants. The bars in the graph represent the Mean ± SD of DESeq2 normalized read counts from RNA-seq of 3 biological replicates. Fold Change Difference: these counts have been additionally normalized using the mean values of the Wild-Type (WT) alone condition. Conditions compared include *DZ2* WT alone, WT + *E. coli*, WT + *B. subtilis*, WT + *C. crescentus*, C10E9 alone and M1 alone.

**FIGURE S7:**
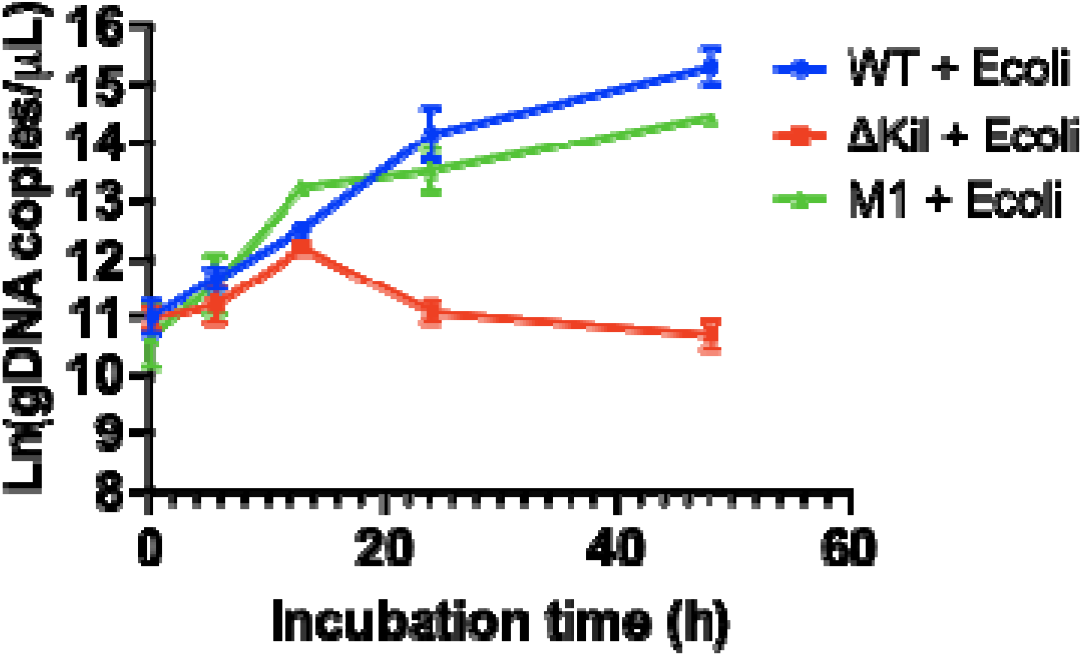
Growth curves of *M. xanthus* DZ2 WT, predation-deficient ΔKil, and evolved mutant M1 during predation on E. coli prey in the presence of 5ug/ml catalase. The scatter plot depicts the natural log-transformed gDNA copies/μL across various predation time points. gDNA concentrations were directly quantified using digital PCR (dPCR). Error bars indicate the SD derived from 3 biological replicates.

**FIGURE S8:**
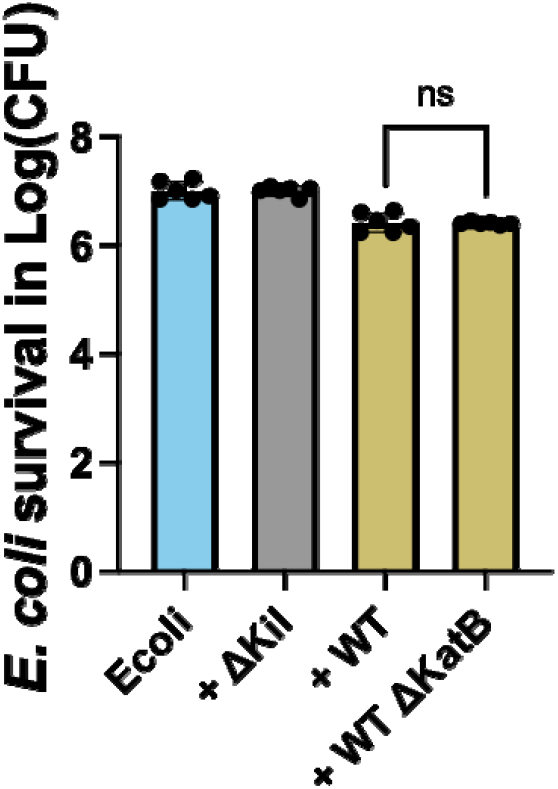
Deletion of *katB* in DZ2 WT strain has no effect on its predation efficiency. The bar graph presents the average survival of *E. coli* MG1655, expressed in log10(CFU), after a 24-hour coincubation with various *M. xanthus* DZ2 predators: WT, predation-deficient ΔKil, and a *katB*-deleted variant. Each bar represents mean +/- SD derived from 3 biological replicates. Statistical significance was determined using one-way ANOVA, with subsequent multiple comparisons conducted using the Bonferroni test; *ns: not significant*.

**FIGURE S9:**
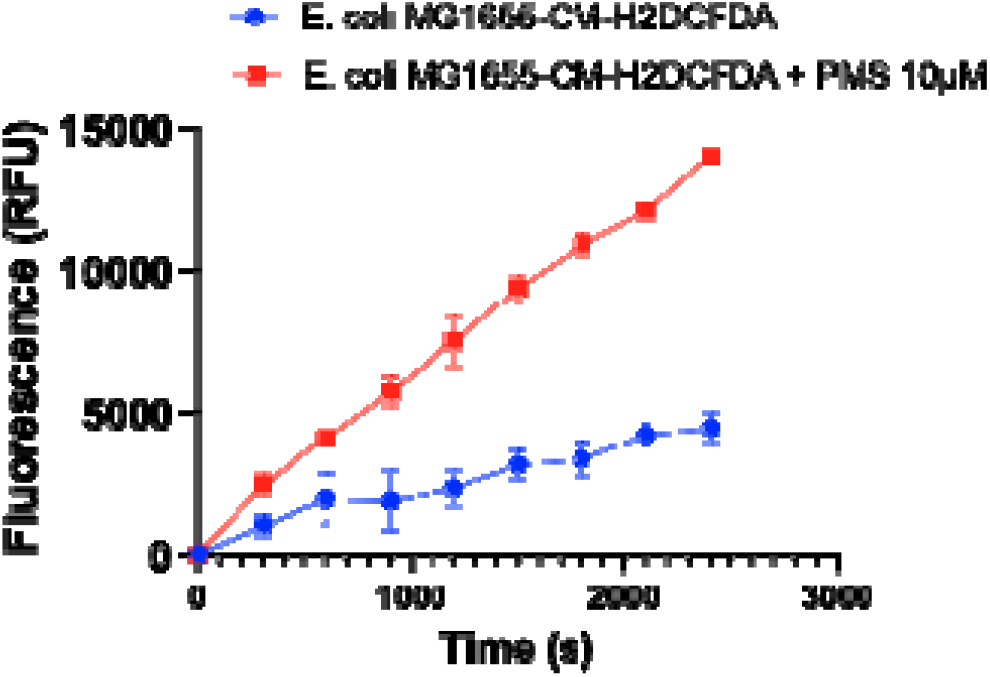
*E. coli* MG1655 cells pre-labeled with CM-H2DCFDA change in fluorescence intensity reflects intracellular accumulation of ROS level in presence of redox-recycling compound phenazine-methosulfate (PMS). Each point represents the mean value of blank-corrected fluorescence intensity +/- SD of three replicates.

**FIGURE S10:**
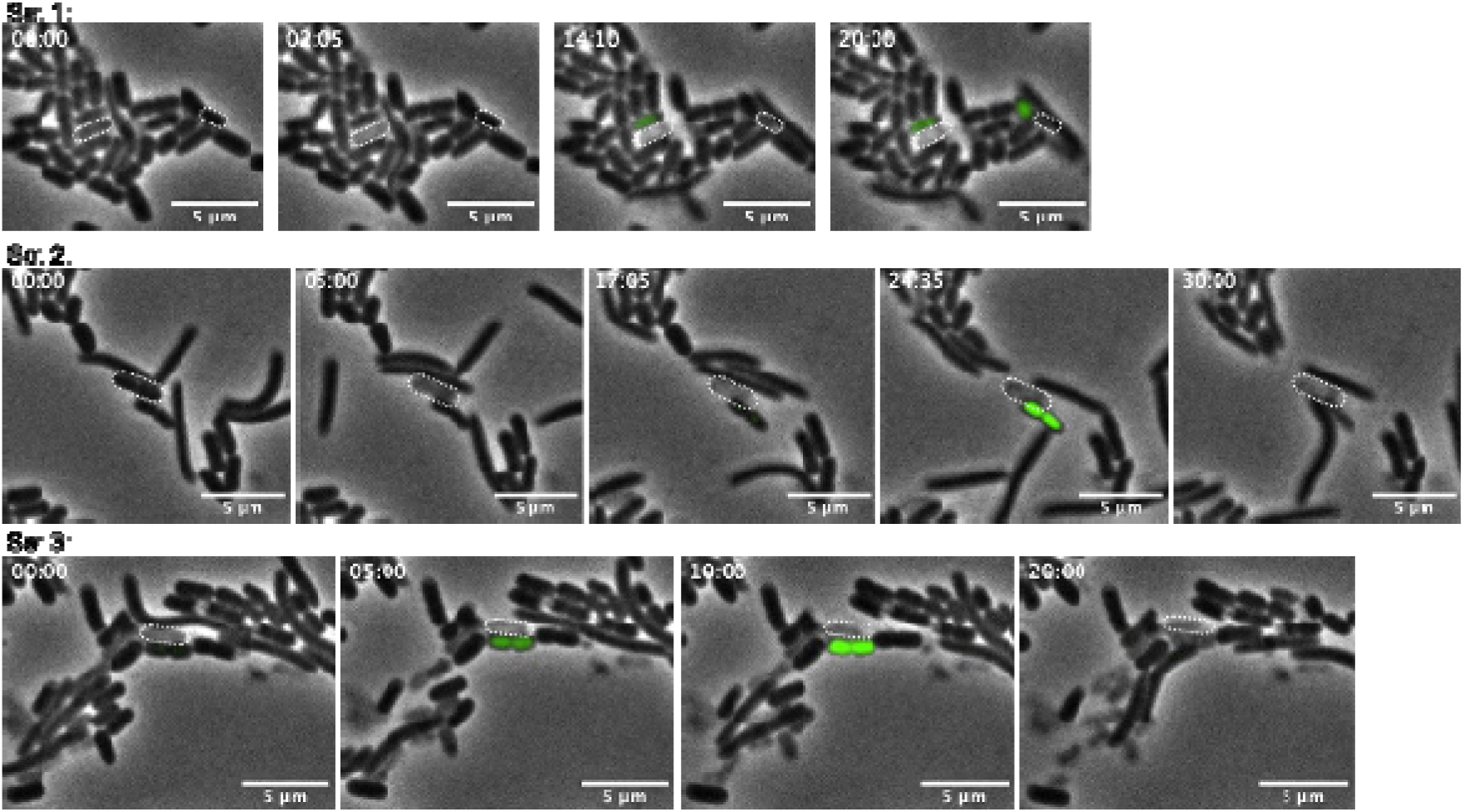
Occasional ROS Burst in Neighboring *E. coli* Cells Induced by PredationTriggered Cell Lysis. Time-lapse fluorescence microscopy was used to study the interaction between *E. coli* prey cells and *M. xanthus* DZ2 WT predator cells. Prior to imaging, *E. coli* cells were stained with the ROS-sensitive dye CM-H2DCFDA. The analysis revealed that in some instances, a ROS burst was observed in *E. coli* cells adjacent to a lysed prey cell. Time points post-interaction with the predator are indicated in the images. Representative images are displayed, selected from two independent experiments.

## SI TABLEs

Table S1: DEGs (a) alone vs *E. coli* (b) alone vs *B. subtilis* (c) alone vs *C. crescentus*

Table S2: 1338 common predatory genes Table S3: DEGs WT vs M1 and WT vs C10E9

Table S4: 441 M1 DEGs similar to common predatome of WT

Table S5: Strains Table S6: Plasmids Table S7: Primers

Table S8: Normalized counts - WT, WT+Ecoli, WT+Bsubtilis, WT+Ccrescentus, C10E9 and M1

Table S9: Annotations and DESeq2 normalized count values for metabolic modules

